# A comprehensive evaluation of CRISPR lineage recorders using TraceQC

**DOI:** 10.1101/2021.10.29.466515

**Authors:** Jingyuan Hu, Hyun-Hwan Jeong, Rami Al-Ouran, Igor Bado, Weijie Zhang, Xiang Zhang, Zhandong Liu

## Abstract

The CRISPR-Cas9 genome editing-based lineage tracing system is emerging as a powerful tool to track cell lineages at unprecedented scale and resolution. However, the complexity of CRISPR-Cas9 induced mutations has raised challenges in lineage reconstruction, which requires a unique computational analysis framework. Meanwhile, multiple distinctive CRISPR-based high-throughput lineage recorders have been developed over the years in which the data analysis is incompatible across platforms. To address these challenges, first, we present the TraceQC, a cross-platform open-source package for data processing and quality evaluation of CRISPR lineage tracing data. Second, by using the TraceQC package, we performed a comprehensive analysis across multiple CRISPR lineage recorders to uncover the speed and distribution of CRISPR-induced mutations. Together, this work provides a computational framework for the CRISPR lineage tracing system that should broadly benefit the design and application of this promising technology.

## Introduction

Determining the origin of cells in multicellular organisms is a long-term goal in developmental biology. Recent advancements in CRISPR-Cas9 genome editing have brought a new generation of lineage tracing techniques that can simultaneously mark cells with irreversible mutations^1-3^. In general, these techniques use CRISPR-Cas9 genome editing to target a specific DNA barcode, which generates diversified mutations that can simultaneously track the cell developmental process. When combined with the state-of-art single-cell sequencing, this lineage tracing technique enables tracking tens of thousands of cells in one experiment. This massive scale lineage tracing has opened up new opportunities to study cell development in single-cell resolution. For example, several studies have applied this technique to map the clonal evolution of cancer metastasis^4-7^.

The CRISPR-Cas9 based lineage tracing technology is a new research field that undergoes rapid expansion. Up to now, several lineage tracing platforms have been developed. GESTALT is one of the first platforms that implemented the idea of CRISPR lineage recording by engineering a target array into the zebrafish’s genome^8^. Using the CRISPR introduced mutations, the GESTALT mapped a complete tissue-level developmental tree of zebrafish. Subsequently, several studies have integrated the CRISPR lineage recorder with single-cell RNA sequencing to provide a cell-level lineage mapping of zebrafish^9-11^ and mouse^12,13^. While most recorders used the canonical CRISPR system, the homing guide RNA (hgRNA) lineage recorder made a major modification by directing the Cas9-hgRNA complex to cuts the DNA locus of itself^14^. This modification has increased the efficiency and diversity of CRISPR genome editing^15,16^.

Across all the CRISPR lineage recorders, the design of the DNA barcodes is vastly different. Some distinctive categories are: 1) Synthesized target array^8,9,12,13^. 2) The hgRNA system^15,16^. 3) Specific DNA sequences that pre-exist in the genome^10,11,17^ (Fig. 1A). Although each platform has different design philosophy on DNA barcode, one commonality is that the barcode sequences are usually compact (less than 300-bp), which makes the high-throughput sequencing readout very homogeneous. As a result, the sequencing analysis could be compatible across multiple CRISPR-based lineage tracing technologies.

**Fig. 1.**
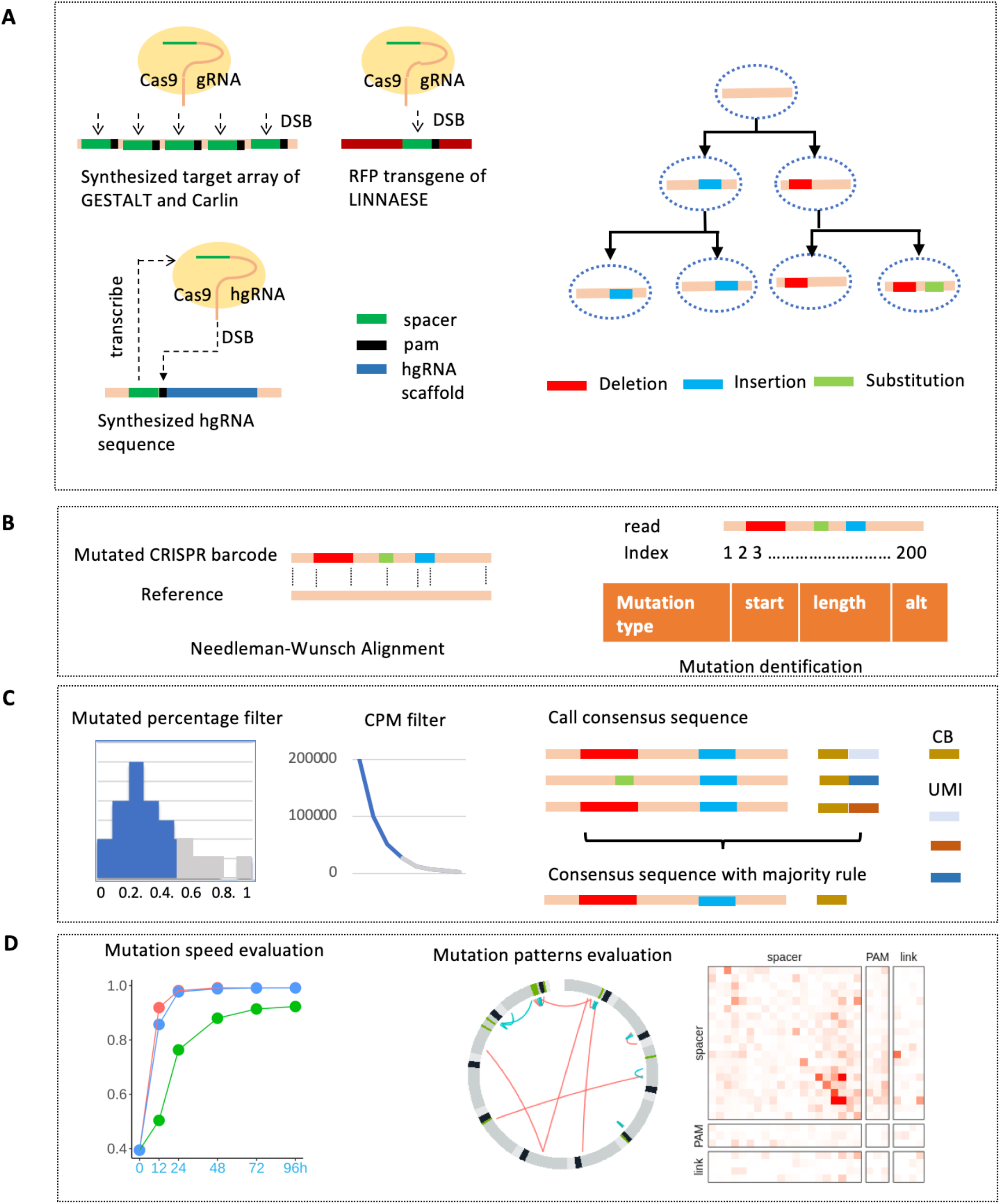
The design of CRISPR lineage recorder and the TraceQC data analysis pipeline. **A** The three distinctive designs of DNA barcodes. 1) The GESTALT and Carlin platform synthesized multiple targets into one barcode, which is transduced into the model organism’s genome. 2) The LINNAESE use CRISPR to edit RFP transgene of an existing zebrafish line *zebrabow M*. 3) The homing guide system use CRISPR to edit the loci of its own guide RNA. The CRISPR-induced mutations allow tracking of cell development. **B** The data processing pipeline of TraceQC. First, Needleman-Wunsch sequencing alignment is performed between the reference and the mutated barcode sequence. Second, the mutated read is index according to the reference sequence. The location information is extracted to represent each mutation. **C** The denoising procedures of TraceQC. First, the contaminated sequences are filtered based on the sequencing alignment score. Next, the sequences with low abundance are removed to denoise the dataset. Finally, for single-cell data, the sequences are collapsed by UMI and cell barcode. **D** Various quality assessments of CRISPR-induced mutations using TraceQC package.

In general, it takes two major steps to reconstruct the lineage tree. Step one is to identify the CRISPR-induced mutations from the sequencing data. A major characteristic of CRISPR-induced mutations is the prevalence of insertions and deletions (indels)^18^. This is primarily a consequence of non-homologous end joining (NHEJ) mechanisms when it repairs the Cas9-induced double-strand break (DSB)^19,20^. To extract indels (ranging from 1-bp to 20-bp) that prevalently exists in the sequencing data, sequencing alignment with affine gap penalty is commonly used. Another characteristic of CRISPR-induced mutations is that the mutation patterns are unevenly distributed. When the Cas9-guide RNA complex cuts the target sequence, the cleavage location is usually a few base pairs upstream to the PAM sequence^21-22^. As a result, the NHEJ repaired DNA sequence is highly deterministic^23^. In fact, the most frequent mutation outcome of the canonical guide RNA system could contribute 10% - 80% of the entire library^24^, which causes a massive scale of parallel evolution (cells from independent lineage acquire the same mutation) of CRISPR barcodes. Therefore, a crucial part of sequencing analysis is to understand the underlying mutation rate caused by NHEJ, which could alleviate the false positivity caused by parallel evolution.

When mutations are extracted from the DNA barcode sequence, step two is reconstructing the lineage tree using mutations. Although classical phylogenetic algorithms such as maximum parsimony^25,26^ are used to reconstruct the large-scale lineage tree^8^, they are not designed to capture the key properties of CRISPR-induced mutations. For example, unlike somatic mutations, CRISPR-induced deletions are typically irreversible. Also, the root of the lineage tree is given by the unmutated sequence, which is usually unobtainable in phylogenetic studies. Since classical phylogenetic algorithms do not consider these characteristics, several studies have designed next-generation lineage reconstruction algorithms using statistical modeling^27-30^, including an open competition^31^.

Although step two is arguably more computationally challenging, a good lineage reconstruction strategy should integrate the two steps seamlessly. However, sequencing analysis is often-times neglected in the lineage reconstruction methodology. For example, LinTIMaT^27^ and GAPML^28^ are two novel reconstruction algorithm that uses maximum likelihood to estimate the lineage tree of GESTALT. Although both methods have considered unique aspects of lineage tree building, such as the importance of mutation rate in parallel evolution, their methodology is purely based on the GESTALT’s mutation metadata. As a result, the connections between the computational model and real experimental data is inexplicit. On the other hand, Cassiopeia^29^, a modified maximum parsimony-based reconstruction methodology, has provided raw sequencing data processing pipeline prior to the reconstruction algorithm. But similarly, the sequencing analysis is secondary in Cassiopeia, in which limited results are shown. We believe a comprehensive sequencing analysis of CRISPR lineage tracing data could benefit these methodologies.

There are several critical challenges in the processing of the CRISPR lineage tracing dataset.1) The CRISPR mutation outcome is highly deterministic, which causes a massive level of parallel evolutions in many lineage recorders. Parallel evolution of barcodes is a significant source of false positives in lineage tracing yet not systematically evaluated in lineage tracing datasets. Research has shown that the CRISPR mutation outcomes could be captured using sequences only, which could be directly used to derive the mutation rate of lineage recorders. 2) The CRISPR editing intensity could significantly influence the lineage recording. For example, intensive CRISPR editing can cause the DNA barcode to saturate too early, preventing the system from recording lineages at a later stage. 3) Since these current methodologies are only performed on one particular dataset, it would be difficult to apply to another CRISPR-based lineage recorder. Moreover, a unified data format for CRISPR tracing data is critically missing, causing the lineage reconstruction strategies incompatible across platforms. Therefore, combing these methodologies with a standardized sequencing analysis pipeline could enable a broader application of the methods across multiple lineage recorders.

Toward this end, we have developed the TraceQC package, which provides a cross-platform data processing pipeline and quality evaluation of multiple sequencing-based CRISPR lineage tracing platforms. Next, by applying the TraceQC package, we have evaluated the key properties of CRISPR mutations across several lineage recorders. Together, the TraceQC package provides a general framework to understand this unique type of data, thereby promoting the application of CRISPR lineage tracing techniques.

## Definitions / Glossary

### Target

the DNA sequence that a Cas9-gRNA complex can edit.

### DNA barcode / barcode

the DNA sequence used to record lineage. It may be composed of a single target or a target array, depending on the specific lineage tracing platform.

### Mutation

an edit in the DNA barcode caused by CRISPR-Cas9.

### Mutation rate

the proportion of a particular mutation in the entire mutation profile. It measures the likelihood of a particular mutation outcome.

### Mutation speed

the number of CRISPR-induced mutations generated in a given period of time. It measures how quick CRISPR edits the DNA barcode. This is referred as rate decay in the GAPML paper.

## Results

### TraceQC identifies CRISPR-induced mutations from sequencing data

The TraceQC pipeline takes raw sequencing data as input and extracts mutations from each sequence (Fig. 1B, Methods). Briefly, Needleman-Wunsch alignment is performed on raw sequences. Due to the nature of the NHEJ DNA repair mechanism, indels are prevalent in CRISPR-induced mutations. Here, we decided to set a small gap opening and extension penalty in the alignment algorithm of TraceQC to favor insertion and deletion over substitution. Next, each aligned sequence is indexed according to the reference barcode in which location information is extracted for each mutation. To reduce technical artifacts (e.g. PCR error, sequencing error, sample contamination), the TraceQC package denoises the dataset using filters based on read count and alignment score (Fig. 1C, Methods). Finally, the package provides a series of functions that evaluate the mutation outcomes across multiple platforms (Fig. 1D, Fig. S13-S28). To illustrate the robustness of the TraceQC package, we analyzed the dataset from all major platforms (Table 1) and revealed differences in mutation characteristics such as speed and rate.

**Table 1.**
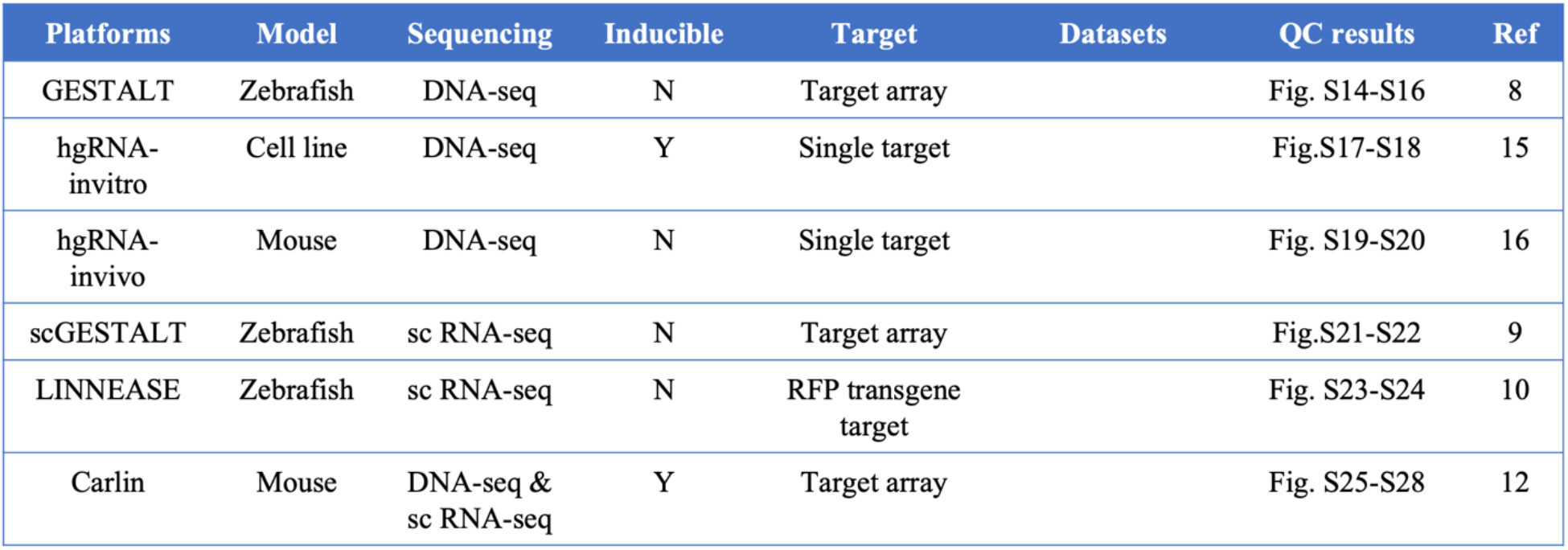
List of CRISPR lineage recorders compared in this study.

| Platforms | Model | Sequencing | Inducible | Target | Datasets | QC results | Ref |
| --- | --- | --- | --- | --- | --- | --- | --- |
| GESTALT | Zebrafish | DNA-seq | N | Target array |  | Fig. S14-S16 | 8 |
| hgRNA-invitro | Cell line | DNA-seq | Y | Single target |  | Fig.S17-S18 | 15 |
| hgRNA-invivo | Mouse | DNA-seq | N | Single target |  | Fig. S19-S20 | 16 |
| scGESTALT | Zebrafish | sc RNA-seq | N | Target array |  | Fig.S21-S22 | 9 |
| LINNEASE | Zebrafish | sc RNA-seq | N | RFP transgene<br>target |  | Fig. S23-S24 | 10 |
| Carlin | Mouse | DNA-seq &<br>sc RNA-seq | Y | Target array |  | Fig. S25-S28 | 12 |

### TraceQC determines preferences of mutation patterns across various platforms

Although most platforms exclude substitutions from the mutation library because they are not as prevalent as indels in CRISPR-induced mutations, it is a viable outcome of NHEJ^32-34^. Using the TraceQC pipeline, we detected the three mutation types: insertion, deletion, and substitution vastly exist in every lineage recorder. On average, three mutations types contribute to a similar diversity to the entire mutation library (Fig. 2A). However, deletion is the most prominent mutation type among the three, as it has the longest length per sequence (Fig. 2B). Each lineage recorder has indel length preference due to differences in experimental designs (Fig. 2C). For example, the barcode sequences of GESTALT and Carlin is a synthesized target array in which deletions can be classified into intra-target deletion and inter-target deletion. While the length of intra-target deletion is mostly less than 10-bp in both GESTALT and Carlin, the length of inter-target deletion could reach 200-bp (Fig S1). In contrast, LINNAEUS is a single-target platform in which CRISPR is used to target red fluoresce protein transgene. Therefore, the targetable region of LINNAEUS has shorter length than target array and we observed that most mutations are concentrated upstream to the PAM sequence (Fig. 2D). Although the homing-guide RNA system is also a single-target platform, its average deletion length is longer than LINNAEUS due to deletions spanning into the extended homing-guide scaffold regions downstream to the PAM sequence (Fig. S9).

**Fig. 2.**
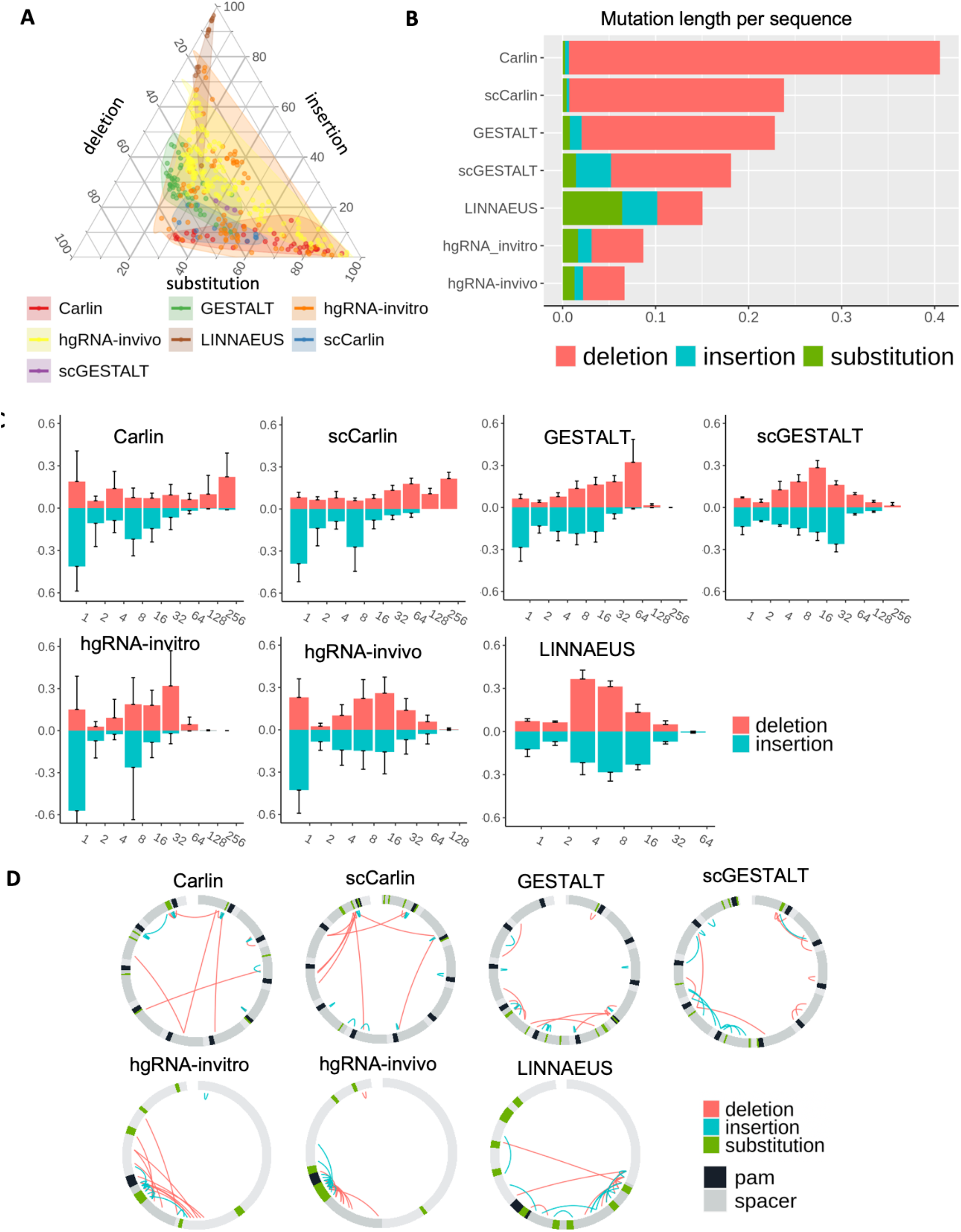
Summary of mutations across multiple CRISPR lineage recorders. **A** The composition percentage of three mutation types: deletion, substitution, and insertion. In the ternary plot, each point is one sequencing sample. **B** The average percentage of mutated length per sequence. The result is averaged across all the samples in each platform. **C** The frequency of indels at a different lengths. The x-axis is the log-scale sequence length by base pair. Each bar aggregate indels length in the range. **D** The top 10 most abundant insertion, deletion, and substitution patterns of each platform. For each platform, the shown mutations are the intersection result of all samples.

Contrary to deletion, the pattern of insertion is more consistent across platforms. In Carlin, GESTALT, and hgRNA system, the most frequent insertion length is 1-bp. Large insertions that are more than 16-bp are rarely seen (Fig. 2C). The mutation properties discovered by the TraceQC are concordant with the original research, and therefore demonstrate that the TraceQC pipeline has provided accurate cross-platform data analysis.

### Impact of mutation speed on the accuracy of lineage reconstruction

Genome editing-based lineage tracing technology requires the accumulation of mutations over time. In current CRISPR lineage recorders, the process of mutation generation is presumably independent of cell division. Therefore, a sound lineage recorder should synchronize the cell division rate with the mutation generations rate such that enough signals are generated to track cell development. Ideally, the mutation speed should be kept moderate. While slow mutation speed will not produce enough signals to mark every cell, quick mutation speed will exhaust the CRISPR barcode too quickly to record a prolonged process (Fig. 3A).

**Fig. 3.**
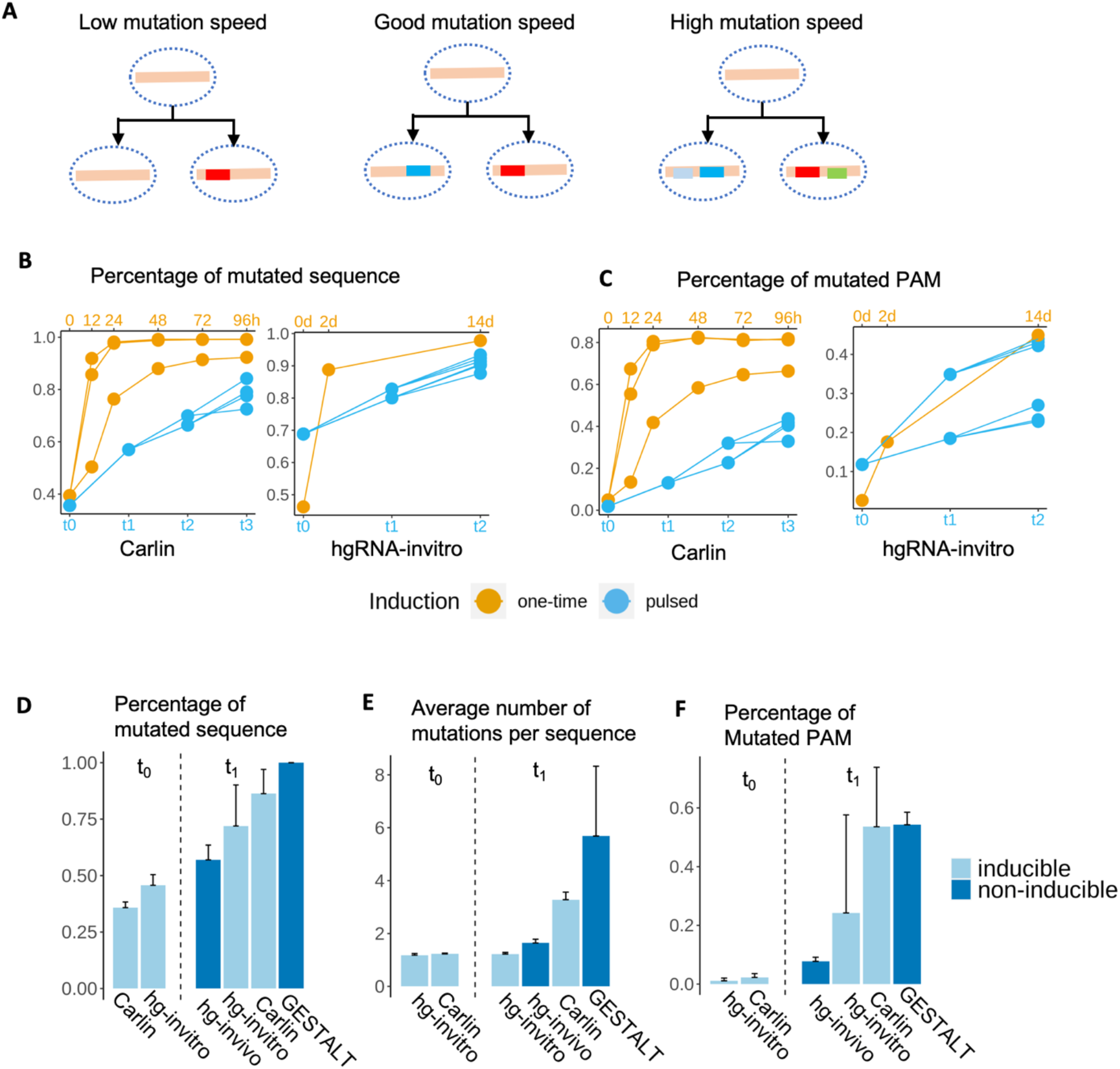
Adjusting mutation speed in CRISPR lineage recorder **A** An illustration of how mutation speed affect lineage recording. While low mutation speed will not produce enough signals for recording, high mutation speed could saturate the barcode. **B** The percentage of the mutated sequence increases through time under two Dox induction methods. The experimental design of Carlin and hgRNA-invitro is described in. In Carlin, the Dox is induced at 0h (one-time), t0, t1, and t2 (pulsed). In hgRNA-invitro, the Dox is induced at 0d (one-time), t_0_, and t_1_ (pulsed). The sample at t0 is sequenced after the Dox induction, which results in a large proportion of mutated barcodes. **C** The percentage of mutated PAM sequence increases through time. **D** The percentage of mutated sequence before (t_0_) and after (t_1_) Dox induction. The non-inducible platforms have constitutional Cas9 expression, which is considered equal as after Dox induction. **E** How many mutations in each sequence before (t_0_) and after (t_1_) Dox induction. **F** What percentage of PAM sequences are mutated before (t_0_) and after (t_1_) Dox induction.

### Adjusting mutation speed in the CRISPR-Cas9 system

Presumably, the mutation speed of CRISPR is determined by the binding efficiency between target sequence and the Cas9 protein. A study has shown that the target sequence, especially the PAM-proximal region is a major determinant of Cas9-target binding efficiency^39^. For example, in the Carlin’s target array, each target can be edited by a perfectly matched guide RNA, which results in similar mutation speed across multiple targets (Fig. S2). However, modifications on the target sequence or sgRNA sequence could decrease mutation speed^39,40^. In the hgRNA-invitro system, researchers engineered the B-21 barcode with multiple PAM binding sites that decreases the mutation speed^41^ (Fig. S3). Besides, the hgRNA system has further synthesized barcodes with various lengths to adjust the mutation speed (Fig. S3 - S4). The result shows the mutation speed becomes slower when there is an increase on the barcode length upstream to the spacer. Although designing a specific target sequence or intentionally creating an off-target event can adjust the mutation speed, these effects are mostly constitutional. Once the target sequences are determined in the system, it is hard to further synchronize them with cell divisions rate.

A more flexible way to control the mutation speed is through the expression level of Cas9 protein. The hgRNA and Carlin system have applied the inducible CRISPR system, whose Cas9 expression is governed by Doxycycline inducible promoter^42^. As a result, the concentration of Dox can positively affect the level of Cas9 expression^43^, which subsequently affects the CRISPR editing activity. In Carlin, researchers have shown that the percentage of mutated barcodes increases in samples with high Doxycycline induction, which indicates an increase of CRISPR activity (Fig. S5).

### Pulsed induction of Doxycycline provides stable mutation speed

When Dox is added at the beginning of the experiment, the consumption of Dox will lead to a drop of Cas9 concentration over time^43^, leading to a drop in the CRISPR mutation activity. Also, a large proportion of the barcode sequence is deleted over time, which preventing the barcode from further editing (Fig. S5). Therefore, the aggregated result is that the mutation speed decreases exponentially through time (Fig. 3B, 3C). Contrarily, a different induction method is to add the Dox several times to replenish the concentration of Cas9 protein, which makes the mutation speed steady (Fig. 3B, C).

### Leaky Cas9 expression introduces background mutations in the inducible CRISPR system

The inducible CRISPR system allows users to initiate the lineage recorder at a desirable time point, which enables lineage tracking of a specific developmental process such as cancer metastasis^5,6^. However, one notable drawback is the leakiness of the inducible CRISPR system, in which Cas9 express before the adding Dox^42,43^. The system leakiness can result in an average of 30%-50% of sequences acquire de novo mutations before the Dox induction. Nevertheless, the system still contains abundant unmutated sequences until it saturates at 90% mutated sequences (Fig. 3D).

Moreover, the barcode sequence could be re-targeted by CRISPR-Cas9, resulting in multiple mutations can appear in one sequence. In hgRNA and Carlin, we found the average deletion length increases after Dox induction, suggesting those barcodes have undergone more than one round of mutation (Fig. 3E, Fig. S6). Besides, the initially mutated sequences have shown limited changes on the PAM sequence (Fig. 3F) and a small percentage of deletions (Fig. S6), which further demonstrated that the sequences could be re-targeted by CRISPR. To sum up, although the leaky Cas9 expression brings background noise to the inducible lineage recorder, the system can still accumulate plenty of mutations subsequently. Therefore, the system’s leakiness will not significantly sacrifice the lineage recorder’s robustness when adjusted correctly.

### The lineage-independent mutation causes parallel evolution of barcodes

In genome-editing-based lineage recorders, parallel evolution is almost inevitable when independent lineage acquires the exact same mutation. This causes cells from independent lineage cannot be distinguishable during tree reconstruction (Fig. 4A). In the CRISPR lineage recorder, parallel evolution is amplified by the highly imbalanced mutation distribution from NHEJ. Besides, the recording capacity of the current lineage recorder usually cannot capture the entire cell population detected by the state-of-art single-cell sequencing, which increases the level of parallel evolution. Therefore, an important goal in lineage tracing sequencing analysis is to uncover the distribution of mutation patterns, thereby improving our understanding of parallel evolutions in these lineage recorders.

**Fig. 4.**
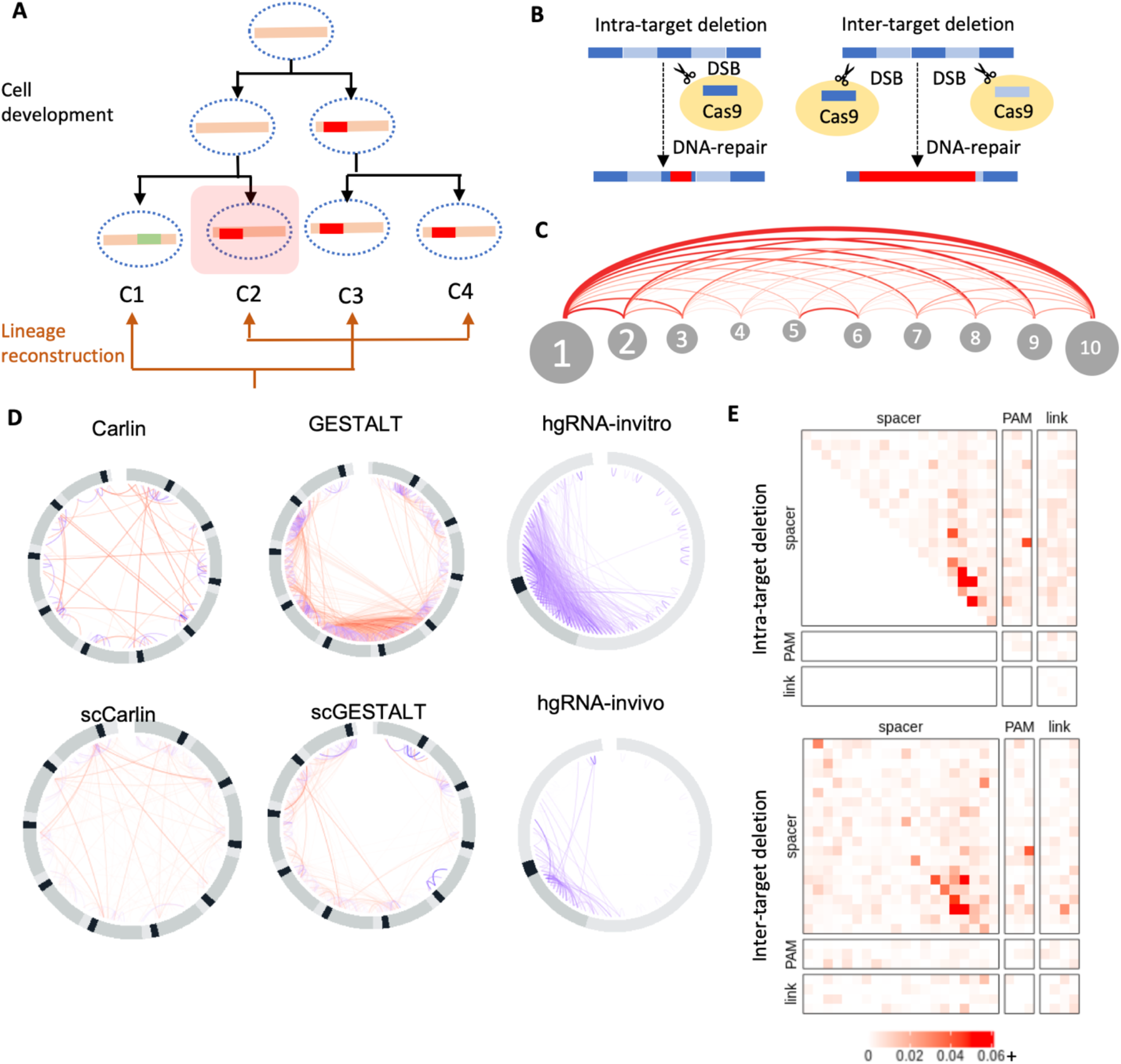
Lineage independent mutations cause parallel evolution in CRISPR lineage recorder. **A** An illustration of parallel evolution leads to undistinguishable lineage assignment. **B** The mechanism of intra-target and inter-target deletion. **C** There are ten targets in the Carlin platform’s target array. The arc diagram shows the frequency of target-to-target interaction of inter-target deletions. **D** The deletion patterns of GESTALT, Carlin, and hgRNA platforms. The red line is inter-target deletion, and the purple line is the intra-target deletion. **E** The position of deletion start and deletion end of Carlin.

### The CRISPR-induced mutations are highly imbalanced

Across all the lineage recorders, we found the most abundant mutation pattern could contribute to up to 60% of the entire mutation library on average, demonstrating a substantial level of parallel evolution of DNA barcodes. Across all the lineage recorders, the CRISPR-induced indels are overwhelmingly concentrated around the PAM sequence (Fig. 4D, Fig S7-S9). This is most likely due to the highly specified DSB location. For GESTALT and the Carlin, the barcode sequences are target array, which means the deletion of both platforms can be classified into inter-target and intra-target deletions (Fig. 4B). The inter-target deletion occurs when two independent Cas9-guide RNA complexes create two DSB simultaneously, which causes drop-out of all the targets in-between (Fig. 4B). In Carlin, the inter-target deletion and intra-target deletion have a very similar mutation hotspot (Fig. 4E, Fig S7).

### Increased mutations diversity improves barcode randomness

In general, increase the diversity of CRISPR mutations can increase the randomness of barcodes^44^, which reduces the number of parallel evolutions. In GESTALT and Carlin, the diversity of barcodes is multiplied by the number of targets in the array. Also, the inter-target deletions further produce additional level of diversity by the number of combinations (Fig. 4C). Similarly, the hgRNA-invivo system uses 60 independent targets throughout the genome to increase the barcode diversity. However, the combinatory effect of multiple barcodes is only detectable via single-cell sequencing. Meanwhile, the hgRNA-invivo system has extended scaffold sequence that allows additional mutations to extend to that region, which generates higher diversity than the canonical guide RNA system (Fig. S9).

Across all the lineage recorders, the three mutation types: deletion, insertion, and substitution have significant differences in mutation distribution. We discovered that insertion and inter-target deletion are more random than substitution and intra-target deletion (Fig. 5A, 5B, Fig. S11A), likely due to a second layer diversity generator.

**Fig 5.**
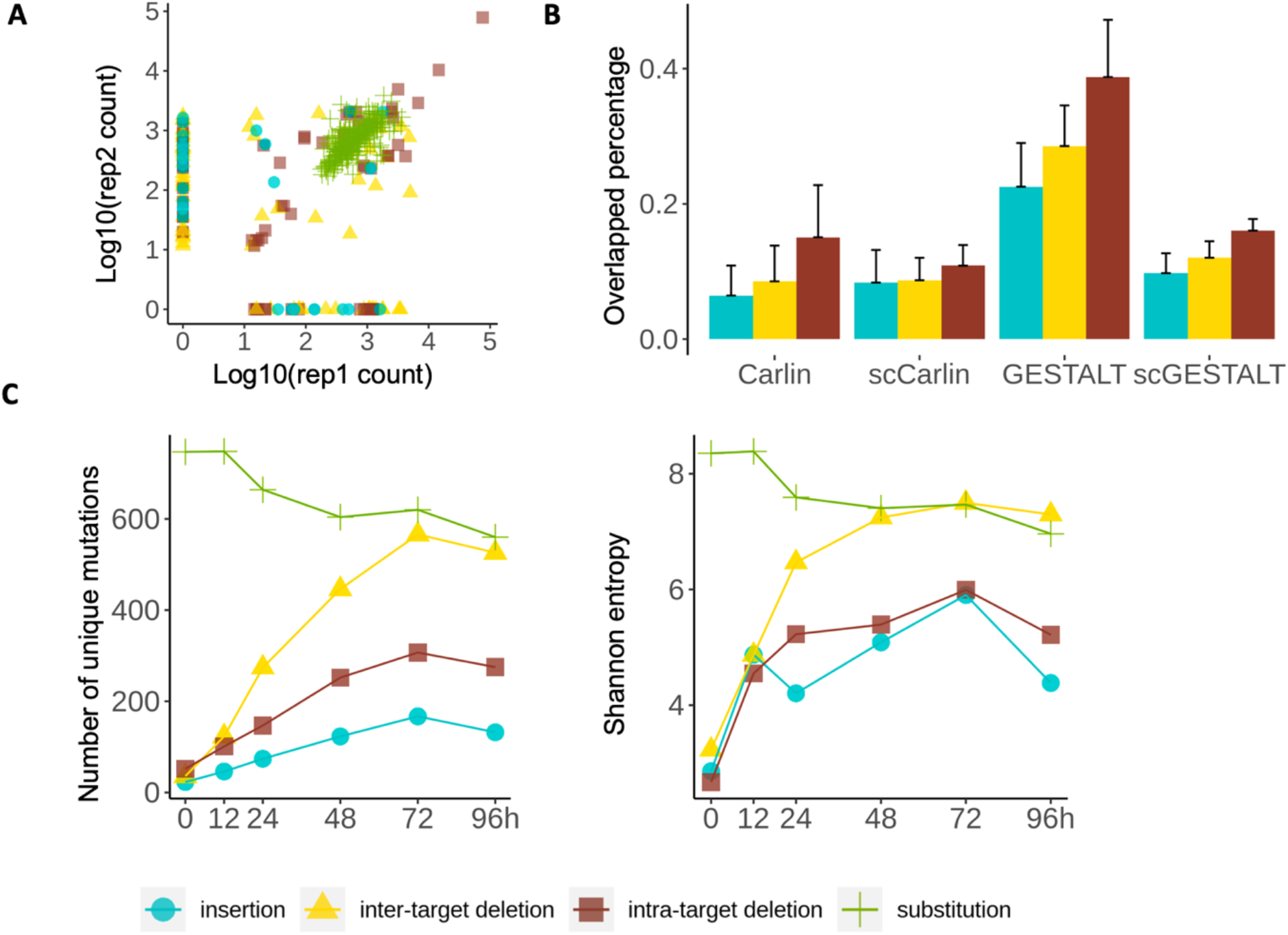
The barcode randomness evaluation. **A** The correlation of replicates of Carlin. The two replicates are negative Dox trial 1 and negative trial 2. The onlapping percentage and Pearson of all mutation types are: inter-target deletion: (19%, 0.61) intra-target deletion: (32%, 0.82) insertion: (14%, -0.14) substitution (100%, 0.61) **B** The percentage of overlapped sequence mutations between independent samples. The independent experiments are selected from different animals, as they have no lineage relations. **C** The change of every mutation type over time. The time-series data is Carlin with a low concentration of Dox induction.

In insertion, the additional diversity is produced by the randomly inserted sequence. In theory, the inserted sequence of length n has a possibility of 4^n^ possible combination, which generates an exponential increase of diversity. As expected, we discovered that the randomness of insertion patterns increases as the length increase (Fig. S10). In GESTALT and Carlin, the increased diversity of inter-target deletion is generated by the random target-to-target interaction. As a result, the inter-target deletion is more random than intra-target deletion (Fig. 5A, 5B).

### Deletion saturates barcode diversity through time

Among all the mutation types, deletion brings the most irreversible damage to the DNA sequence. When evaluating the accumulation of each mutation type over time, we discovered that inter-target deletion had gained the most diversity whereas the diversity of insertions and substitutions are gained little or decreased (Fig. 5C, Fig. S11). This is mainly because the DNA barcode is gradually saturated by deletion over time as the prolonged CRISPR activity removes a great proportion of the barcode sequence (Fig. S6). As a result, the barcode is too exhausted to bear substitutions and insertions in later development stage. In addition, in GESTALT and Carlin, the inter-target deletion removes all targets in between, which completely clean the diversity of a large proportion of barcode. Contrarily, substitution and insertion have limited diversity when the barcode. In contrast to Carlin, the hgRNA is a single-target platform in which the deletions are shorter. Therefore, the domination of deletion in the mutation library is mitigated compared to GESTALT and Carlin (Fig. S12).

## Discussion

The two properties of CRISPR-induced mutations: speed and rate, have a broad impact on lineage reconstruction. In the non-inducible CRISPR system, the mutation speed depends entirely on the binding between the Cas9 and the target sequence. But in the inducible CRISPR system, the mutation speed can also be controlled by Dox concentration, bringing flexibility and accessibility to the recorder. Moreover, the pulsed induction of Dox enables a stable expression of Cas9, which results in a steady CRISPR mutation speed. The current application of CRISPR lineage recorder usually focuses on a specific developmental process in which the cell division rate is steady. Therefore, pulsed induction of Dox could be useful in many experiments’ settings. However, in many organisms, the mutation rate of cell division varies among different tissues and different developmental stages, e.g. mouse embryogenesis^45^. Also, studies have shown that CRISPR activity varies in different tissues^46^. This could bring challenges to adjusting speed in lineage recorders. Moreover, the current pulsed induction experiment in Carin and hgRNA is only performed in in-vitro systems. In in-vivo experiment, closely monitoring the Cas9 expression level could be challenging. Determining the optimal Dox concentration prior to the in-vivo experiment to can help identifying the desired mutation speed.

The randomness of CRISPR mutations affects the level of parallel evolutions in lineage recorders. In general, increase the diversity of barcodes by incorporating multiple targets could greatly increase the barcode randomness. We demonstrated that the inter-target interaction of the GESTALT and Carlin platform generated increased the mutation diversity, thus provide robust lineage barcoding through time. The hgRNA mouse model used 60 independent targets throughout the genome to increase the barcode diversity. However, the current experiment readout by bulk DNA sequencing cannot detect the independent target interactions, which could be evaluating using single-cell sequencing.

Meanwhile, in GESTALT and Carlin, we discovered that inter-target deletion might saturate the DNA sequence too quickly, preventing insertion and substitution from occurring. Moreover, inter-target deletion could dropout existing mutations on the sequence, thereby reduce the barcode diversity. Therefore, when building the lineage tree in GESTALT and Carlin, it is important to modeling the dropout effect of mutations.

Potentially, reduce the number of deletions could increase the duration of CRISPR lineage recorders. Recently, many variants of CRISPR-based genome editing technologies have been developed. For example, the base editor^49-52^ brings a single nucleotide change to the genome that could provide sustainability to the barcode.

One limitation of our mutation distribution analysis is the lack of quantitative measures. Although several studies have demonstrated that NHEJ-induced mutation patterns can be predicted in-silico^24,35-38^, these machine learning models cannot be directly applied to the CRISPR lineage tracing dataset. First, the NHEJ mechanism varies among different organisms, which results in different mutation patterns^36^. Second, these models are trained on a single-target canonical guide RNA library. It cannot capture the properties of target array or hgRNA system. Nevertheless, from the lineage tracing dataset, we discovered the characteristics of CRISPR-induced mutations are mostly concordant with the previously identified characteristics, such as the prevalence of 1-bp insertion. More importantly, we found the mutation frequencies are highly correlated between replicate, which means a machine learning solution could be applied to predict the mutation distribution.

The scale and complexity of single-cell lineage tree bring challenges to the classical phylogenetic algorithms^50,51^. Also, the CRISPR-induced mutations require a novel computational modeling framework. Unsurprisingly, every new lineage reconstruction algorithm has realized the importance of mutation rate. However, directly estimating the mutation rate from the lineage tracing data is one of the biggest challenges, which causes them to creatively detoured the problem. For example, the GAMPL algorithm uses Markov chain to model mutation generation. Instead of estimating the mutation rate, GAMPL used lumped matrix to estimate the transition rate between meta-states. In contrast, Cassiopeia obtained the mutation rate through experiment. They performed an in-vitro experiment of CRISPR barcoding for 15 cell cycles. Then mutation rates are derived empirically using frequency. Although TraceQC does not directly provide an estimation of mutation rate, we believe it could be obtained from a combination of *in-vitro* experiment and *in-silico* mutation frequency prediction. Next, an integration between the sequencing analysis pipeline and tree reconstruction algorithms could produce a more accurate lineage reconstruction strategy.

Overall, the single-cell resolution CRISPR lineage tracing is a promising technology with many potential applications. We developed the TraceQC package to provided sequencing analysis and quality evaluation of lineage tracing datasets across multiple platforms. We hope our study could facilitate a wider application of this technology and provide some insights into its future development.

## Methods

### TraceQC pipeline

The TraceQC R package is available at https://github.com/LiuzLab/TraceQC. The tutorial of TraceQC is available at https://github.com/LiuzLab/TraceQC/wiki. The input of TraceQC pipeline are raw sequencing files and annotated barcode construct sequence. The workflow is shown in Fig. S13. The output is CRISPR-induced mutations and various plots of quality evaluation.

### Sequencing alignment

The first step of TraceQC is to align reads to the reference sequence. The reference sequence must be annotated with a CRISPR targetable region because the raw reads typically contain adapter sequence. The Needleman-Wunch sequencing alignment with affine gap penalty is applied to each raw sequence. This alignment requires the user to provide four parameters: match, mismatch, gap open penalty, and gap extension penalty. The default parameters are: match = 2, mismatch = -2, gap open = -6, gap extension = -0.1. After the sequencing alignment, the CRISPR targetable region is selected for the next step.

### Alignment quality evaluation

The alignment quality filter is used to remove reads that are contaminated. In every CRISPR lineage recorder, the barcode sequence is usually amplified and sequenced specifically. However, some samples could contain up to 20-40% reads that do not come from the CRISPR barcode. Using the alignment score difference, it is usually easy to separate the contaminated sequence from the mutated CRISPR barcode (Fig. S14C). To quantitively determine the contaminated sequence, a decoy sequence library is generated by randomly substitute ε percentage of the sequence. Next, TraceQC trains a local regression model between the substituted percentage and the alignment score. Finally, the sequence below the optimal substitute percentage can be selected.

### Identify CRISPR-induced mutations

After the sequencing alignment, the CRISPR targetable region of each sequence is extracted. Next, TraceQC scans through the targetable region of each sequence and extracts every mutation. Basically, TraceQC extracts four properties for each mutation: mutation type (insertion, deletion, and substitution), starting position, indel length, altered sequence. Finally, the TraceQC summarizes every mutation into a count table.

### Read merging

In single-cell RNA sequencing, a unique molecular identifier (UMI) is used to distinguish unique mRNA transcripts. The cell barcode is used to determine cell identity. Presumably, when the CRISPR barcode is transcribed, the mRNA transcript from each cell should be identical. Therefore, TraceQC provides merging functions for mutations in each cell. First, for each cell, reads with the same UMI are grouped together. The mutations that appears in more than 50% of reads are retained. During this step, UMI with a low read count should be filtered according to the guideline of the particular single-cell sequencing platform. Next, for each UMI in each cell, a second merger is applied to retain mutations that appear in more than 50% UMI.

### QC plots

There are three main aspects that TraceQC evaluates: 1) The quality of sequencing alignment. Using alignment score, TraceQC removes the percentage of contaminated sequences. The method is described in the section: *alignment quality evaluation*. 2) Mutation patterns can be evaluated from the circular plot. Users can visualize the position of each mutation and identify the mutation hotspots. 3) For single-cell lineage recorders. TraceQC determines the read count distribution of UMI, which allows users to filter. Also, the mutation distribution across cells allows users to understand the lineage recording ability.

### Processing of bulk DNA-seq datasets

In this study, we analyzed bulk DNA-seq datasets from Carlin, GESTALT, hgRNA-invitro, and hgRNA-invivo platforms. The bulk DNA-seq dataset of Carlin used paired-end sequencing. We merged read R1 and R2 using software FLASH with default parameters. Next, the dataset is processed as follows:

1. The merged read is processed using the TraceQC alignment with the default parameter. We used ε = 0.4 to filter out contaminated sequences.
2. We normalized each sequencing sample using count per million (CPM). Reads with CPM > 10 are retained.
3. CRISPR-induced mutations are identified using TraceQC.

We processed a part of GESTALT’s in-vivo bulk DNA-seq dataset (GESTALT paper fig. 3). Although both GESTALT and Carlin used paired-end sequencing, the reads of GESTALT cover approximately 60% of the entire barcode, whereas Carlin covers 100%. Therefore, we processed the R1 and R2 of GESTALT separately using the same procedure. Finally, the mutation event of R1 and R2 are merged for each sample.

The hgRNA-invitro dataset used single-end sequencing. The complete dataset is processed using the same procedure as Carlin.

The hgRNA-invivo platform contains 60 independent CRISPR barcodes. A DNA identifier is assigned to each barcode which presumably cannot be edited by CRISPR. Therefore, we first used the 12 base pairs DNA sequence upstream to locate the DNA identifier. In this step, the Levenshtein distance of 1 is allowed for the 12-bp sequence. Next, the 10-bp directly downstream is extracted to match with the identifier. The sequences are grouped by each identifier and processed by the TraceQC pipeline using the same procedure as Carlin.

### Processing of single-cell RNA-seq datasets

In this study, we analyzed single-cell RNA-seq datasets from Carlin, LINNAEUS, and scGESTALT. The single-cell dataset of Carlin used 10X Chromium sequencing. First, the raw sequences of CRISPR barcodes are processed with cell ranger V3.1, in which the error-corrected cell barcodes and UMI are identified. Next, the dataset is processed as follows:

1. The merged read is processed using the TraceQC alignment with the default parameter. We used ε = 0.4 to filter out contaminated sequences.
2. For each cell, reads are grouped by UMI. UMIs with read count < 10 are filtered out.
3. CRISPR-induced mutations are identified for each read using TraceQC.
4. To merge reads in each cell. First, for each UMI, mutations appear in more than 50% reads are retained. Second, for each cell, mutations that appear in more than 50% UMI is retained.

The LINNAEUS also uses 10X Chromium sequencing. First, the raw sequences of CRISPR barcodes are processed with cell ranger V2.0.2, in which the error-corrected cell barcodes and UMI are identified. However, the read length of 10X single-cell RNA-seq cannot cover the entire barcode region. Therefore, we performed semi-global sequencing alignment that does not penalize the end gap. The other processing pipeline is the same as single-cell Carlin.

The scGESTALT uses in-Drops single-cell RNA-seq. It applies the v7 barcode sequence of GESTALT. According to the annotation, we first used the 10 base pairs DNA sequence downstream to extract the UMI. The rest procedure is the same as single-cell Carlin.

### Mutation properties analysis

Using the TraceQC identified mutations, we first calculated the number of unique mutations per sample. Each unique mutation is characterized by mutation type (insertion, deletion or substitution), starting index of the mutation according to the reference sequence, mutation length and altered sequence (only for insertion and substitution). The ternary plot shows the relative ratio of unique mutations in each type.

### Time-series data analysis

Across all the datasets we analyzed, only Carlin and hgRNA-invitro have time-series data. Briefly, the Carlin platform used CRISPR to target the embryonic stem cells (ESCs) of mouse. In the one-time induction experiment, the ESCs were exposed with low (0.04 μg/ml), medium (0.20 μg/ml) and high (1.0 μg/ml) of Dox. Then, bulk DNA sequencing is performed at 0 hours (before Dox induction), 12-hours, 24-hours, 48-hours, 72-hours and 96-hours. In the pulsed-induction experiment of Carlin, the ESCs is exposed to three pulses of Dox (0.04 μg/ml) every 6 hours. After each exposure, cells were picked for sequencing and further outgrowth.

The experiment design of hgRNA-invitro is similar. In the one-time induction experiment, the 293T human cell line was exposed to Dox (2.0 μg/ml). Then, bulk DNA sequencing is performed at 0 days (before Dox induction), 2 days, and 14 days. In the pulsed-induction experiment of hgRNA, the 293T cells were first exposed to two hours of Dox to create the founder clone. Next, the Dox is removed while the founder clone grew into 6 clones. To further expand the cell populations, 100 cells are selected for further expansion by exposing them to Dox for two hours. In our study, we compared the mutation speed of one-time induction and pulsed induction of A21 barcode (Fig. 3B, 3C).

As for mutation speed analysis, we calculated the percentage of the unmutated sequence out of each sample. Next, we locate the PAM (NGG) sequence of both platforms. Next, we considered the PAM is mutated when either of the two guanines is mutated. When calculating the percentage of PAM for Carlin, all 10 PAMs are treated equally.

### Mutation hotspot analysis

For GESTALT and Carlin target array, the construct sequence designs are similar in which each target (contains spacer and PAM) is separated by a short linker sequence. For both platforms, we defined each target as each consecutive sequence that contains spacer, PAM, and linker. To classify deletion into intra-target deletion and inter-target deletion, we simply determined if the starting position and ending position belong to the same target. As for the mutation position analysis, the heatmap and histogram show the average result of all samples.

### Mutation dependency analysis

In the mutation dependency analysis, first, we selected an independent experiment of Carlin and hgRNA-invitro before induction because the CRISPR mutations in these samples are least confounded by cell development. Next, for Carlin, scCarlin, GESTALT, scGESTALT, we calculated the mutation dependency between samples that were not taken from the same animal/cell populations.

## Data and code availability

All raw sequencing data in this study is publicly available and can be downloaded from GEO. The corresponding GEO accession number is listed in Table 1. The TraceQC R package is available in GitHub (https://github.com/LiuzLab/TraceQC) under the MIT license. The code for regenerate the analysis results of this manuscript is available at https://github.com/LiuzLab/TraceQC-manuscript.

