## Supplemental figures for "A comprehensive evaluation of CRISPR lineage recorders using TraceQC"

**A**

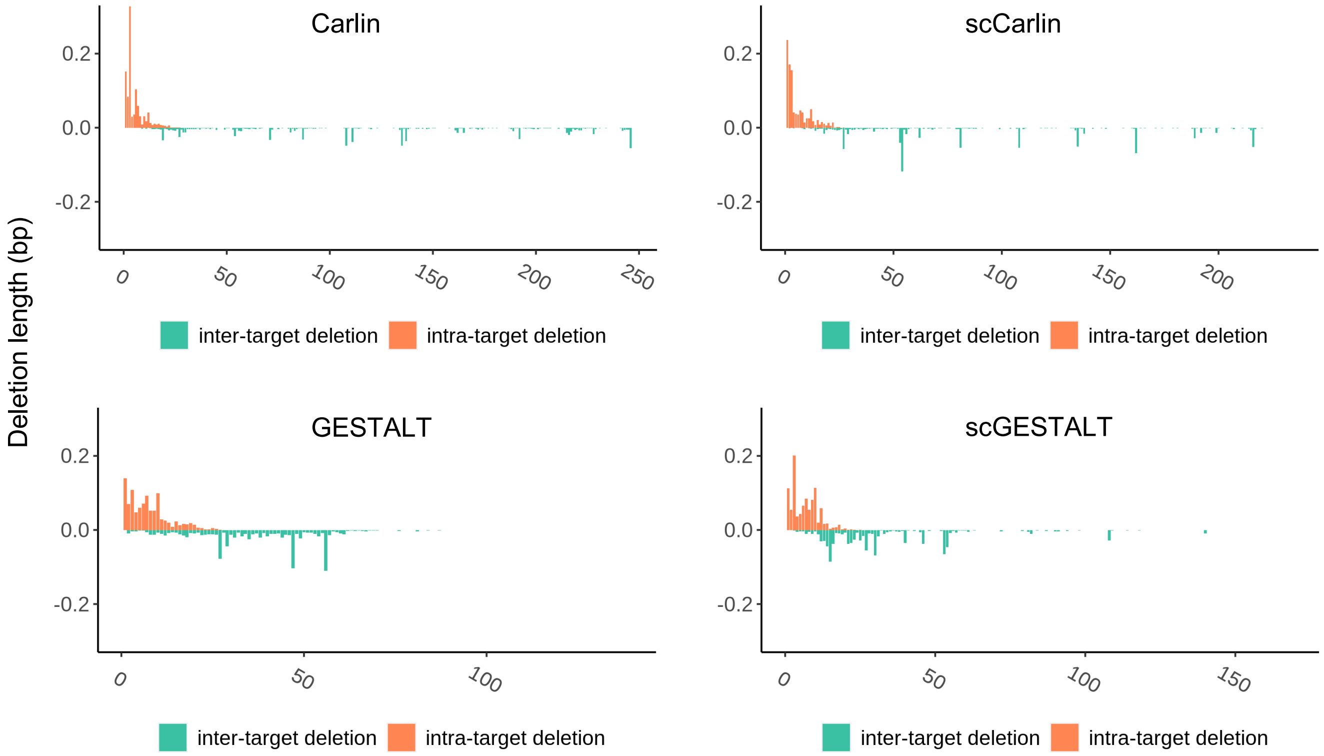


**Figure S1. A** The deletion length (bp) of intra-target deletion and inter-target deletion.

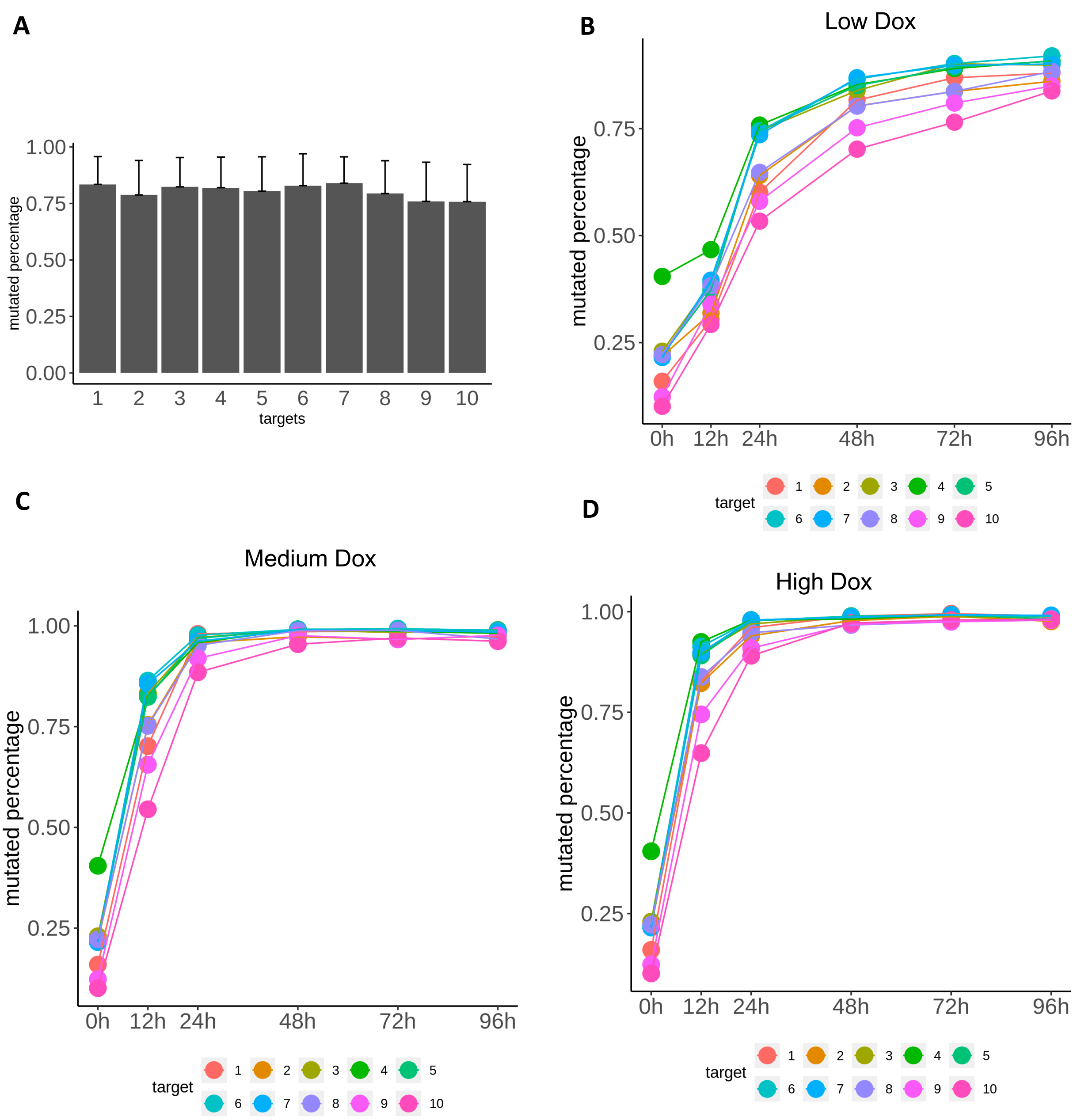


**Figure S2.** The mutation speed of Carlin by each target. **A** The percentage of mutated targets at 96h. **B - D** The change of mutated targets under different concentration of Dox.

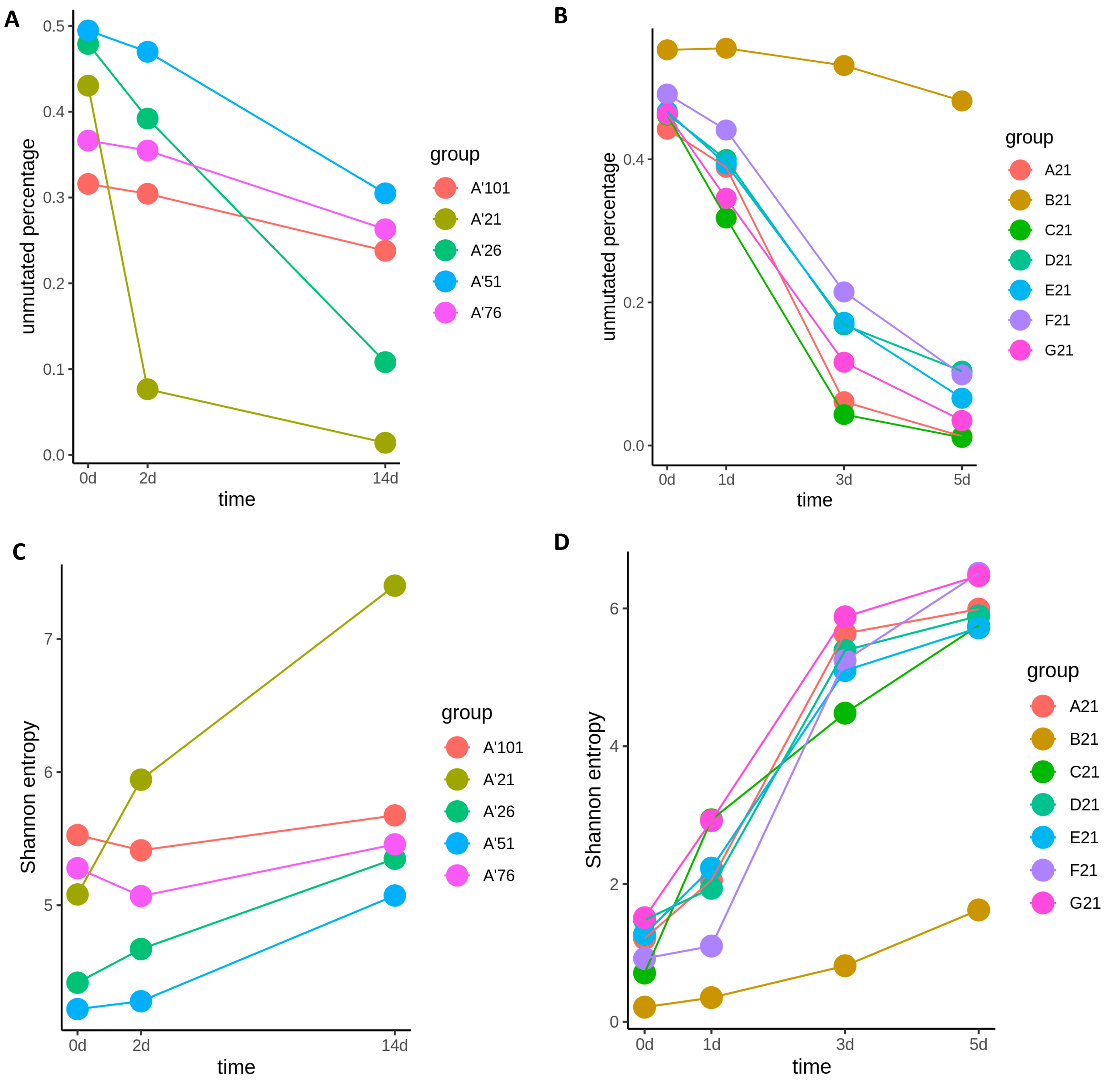


**Figure S3**. The time-series data of hgRNA-invitro . **A-B** The change of mutated sequence percentage through time. The hgRNA-invitro have designed multiple barcodes with different length, e.g. the spacer and pam makes up 21 base pairs in A21 barcode. **C-D** The change of Shannon Entropy through time, which is calculated using indels only.

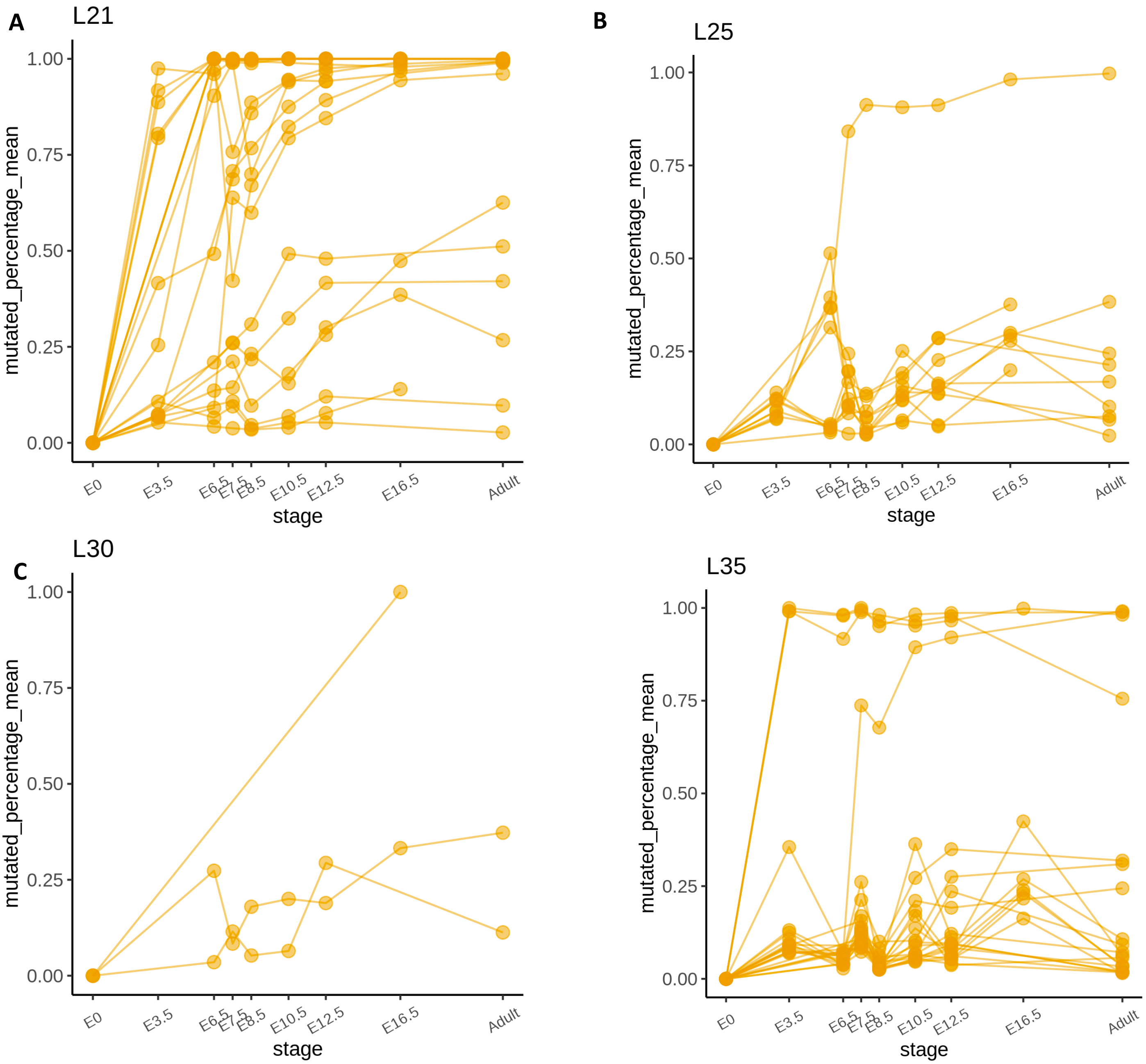


**Figure S4**. The time-series data of hgRNA-invivo. **A - D**. The percentage of mutated barcode with different length. The mutated sequences are considered with indels only.

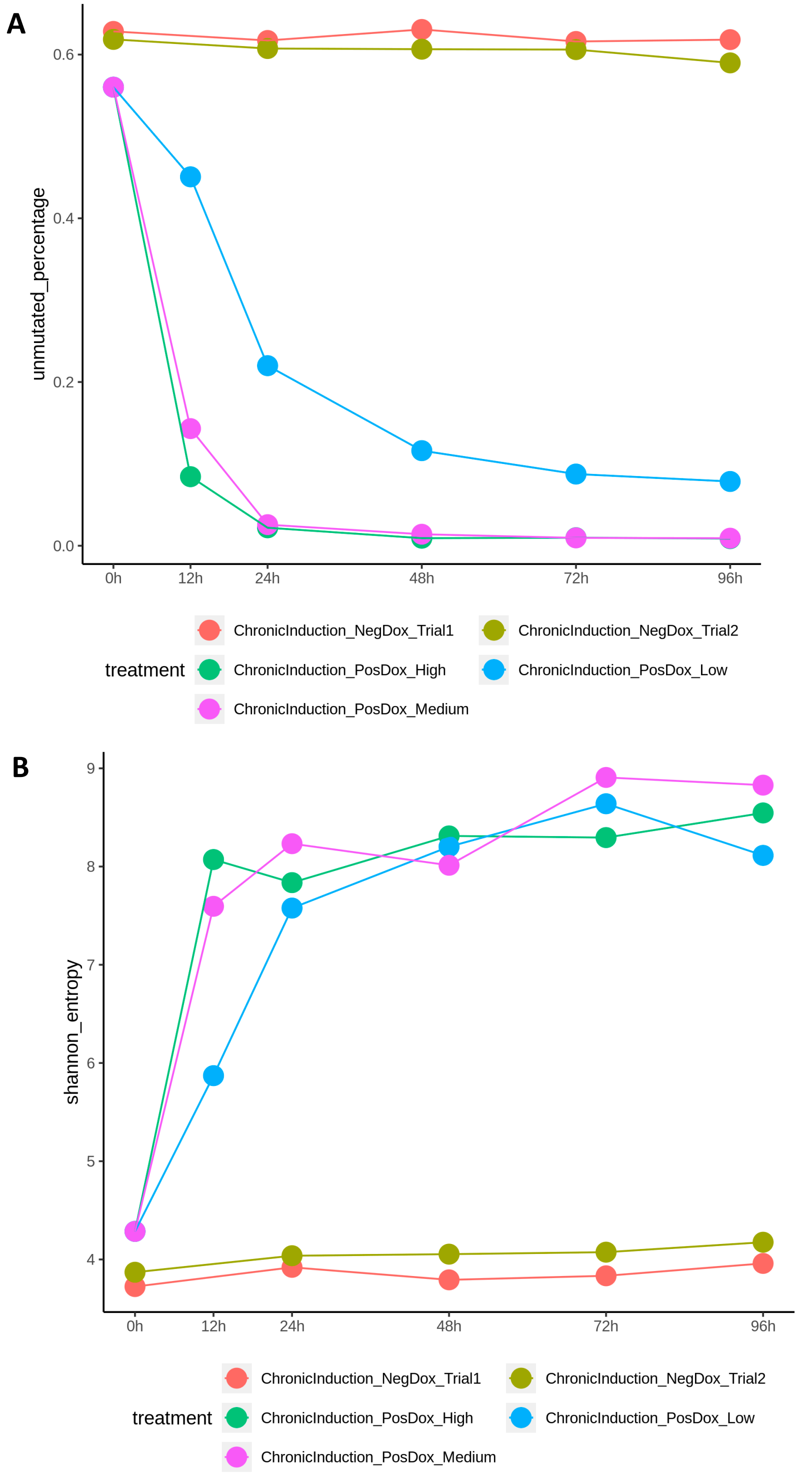


**Figure S5**. The time-series data of Carlin under low, medium and high concentration of Doxycycline induction. **A** The change of unmutated sequence percentage through time. **B** The change of Shannon entropy through time.

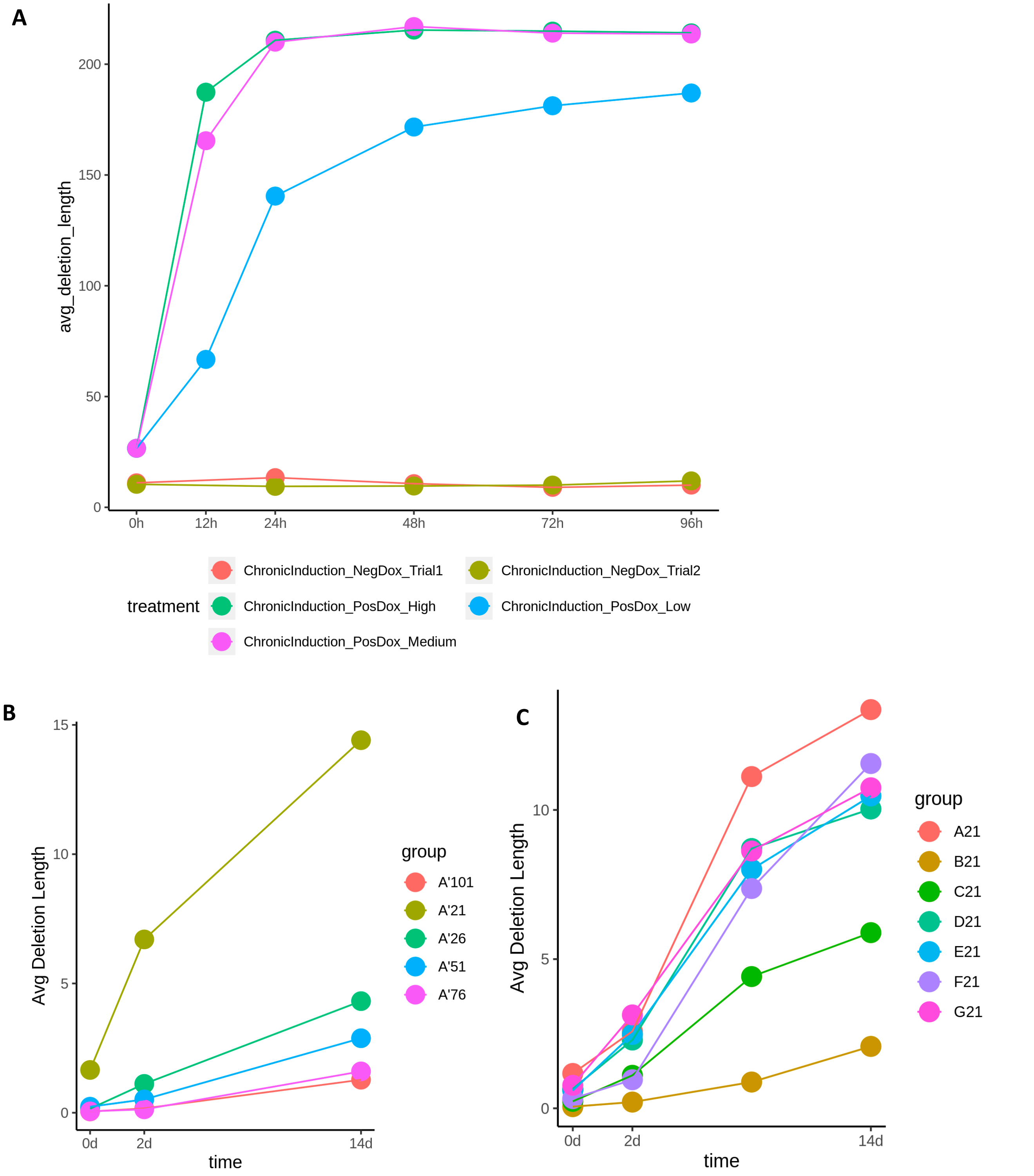


**Figure S6**. The change of deletion length (bp) through time. **A** The average deletion length of Carlin's time-series data. **B - C** The average deletion length of hgRNA-invitro's time-series data.

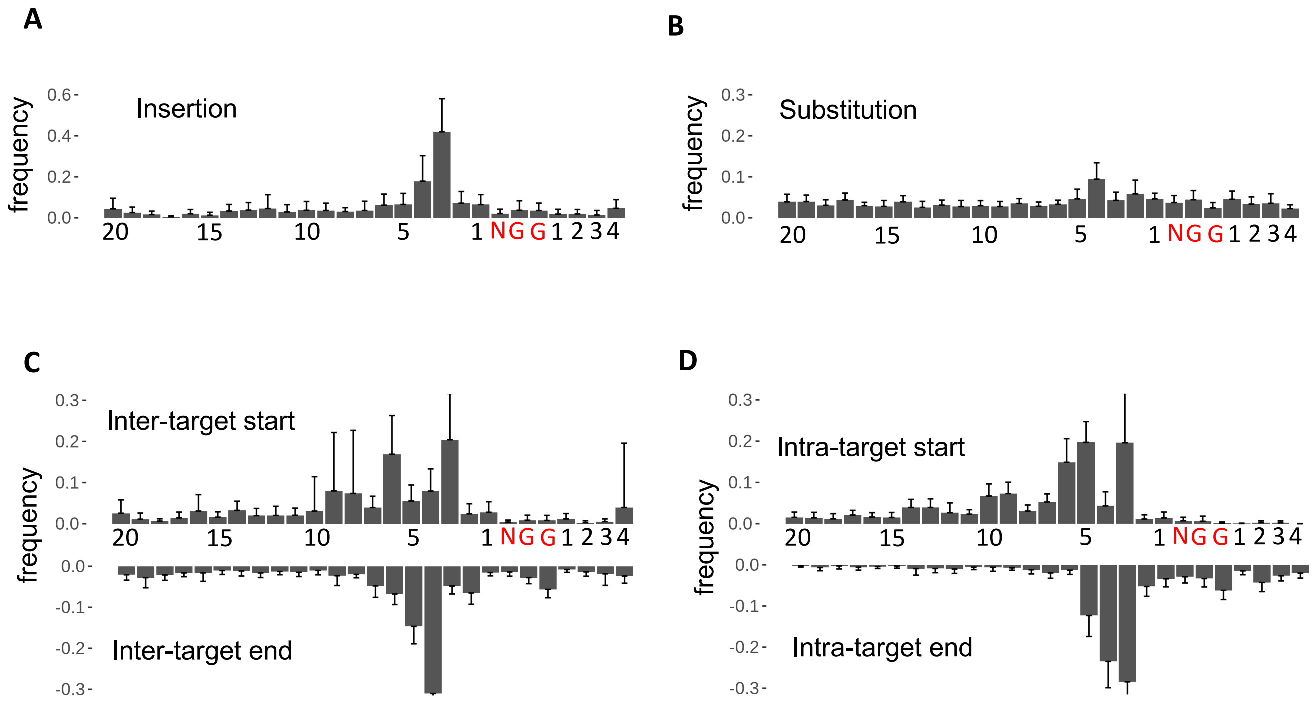


**Figure S7**. The histogram of Carlin's mutation position. The ten targets of Carlin are aggregated together. **A** Insertion. **B** Substitution. **C** Inter-target deletion. **D** Intra-target deletion.

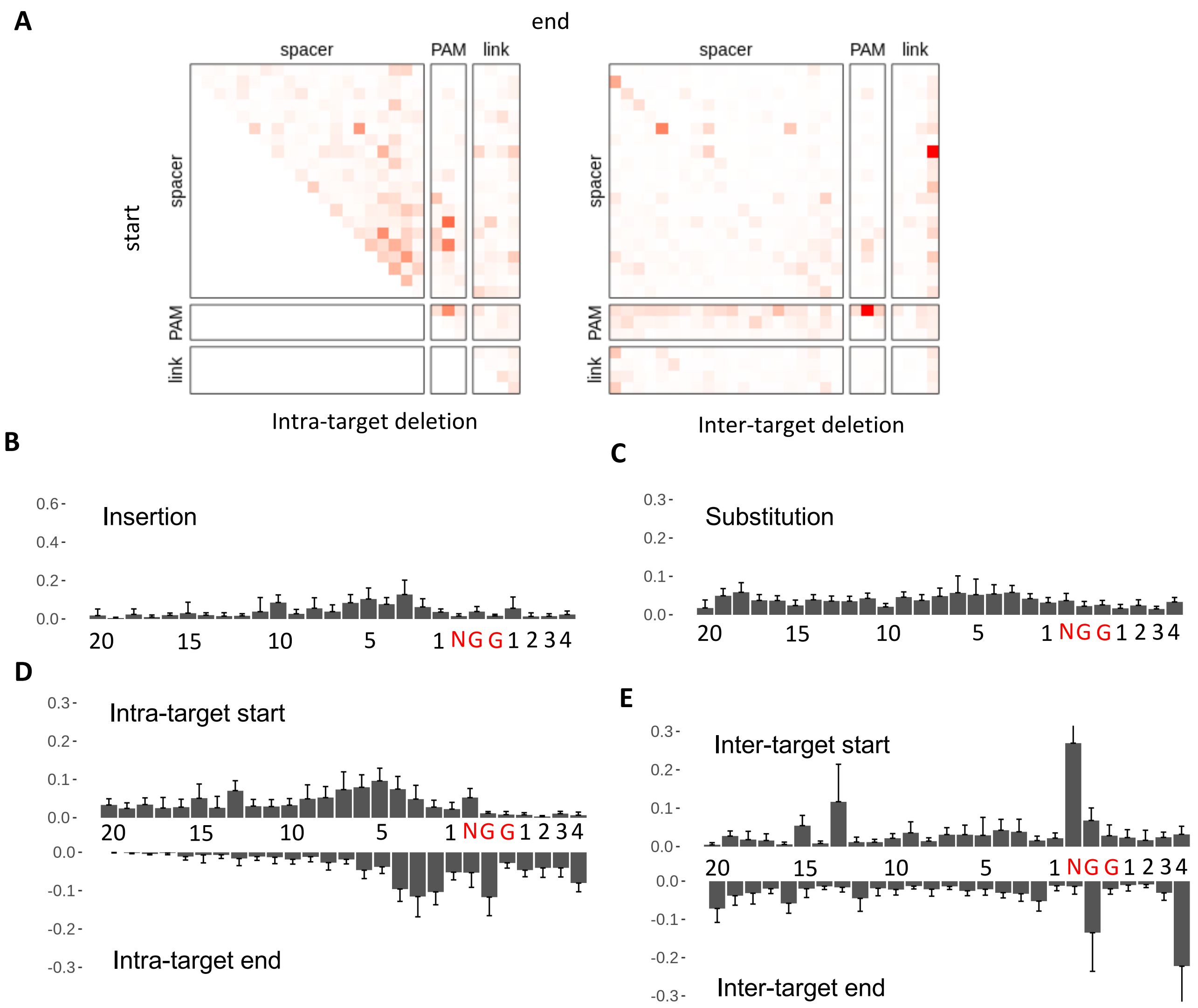


**Figure S8** The frequency of GESTALT's mutation position. The 10 targets of GESTALT are aggregated together. **A** The heatmap showing the start and end of intra-target deletion and inter-target deletion. **B** The histogram of GESTALT's insertion position. **C** Substitution. **D** Inter-target deletion. **E** Intra-target deletion.

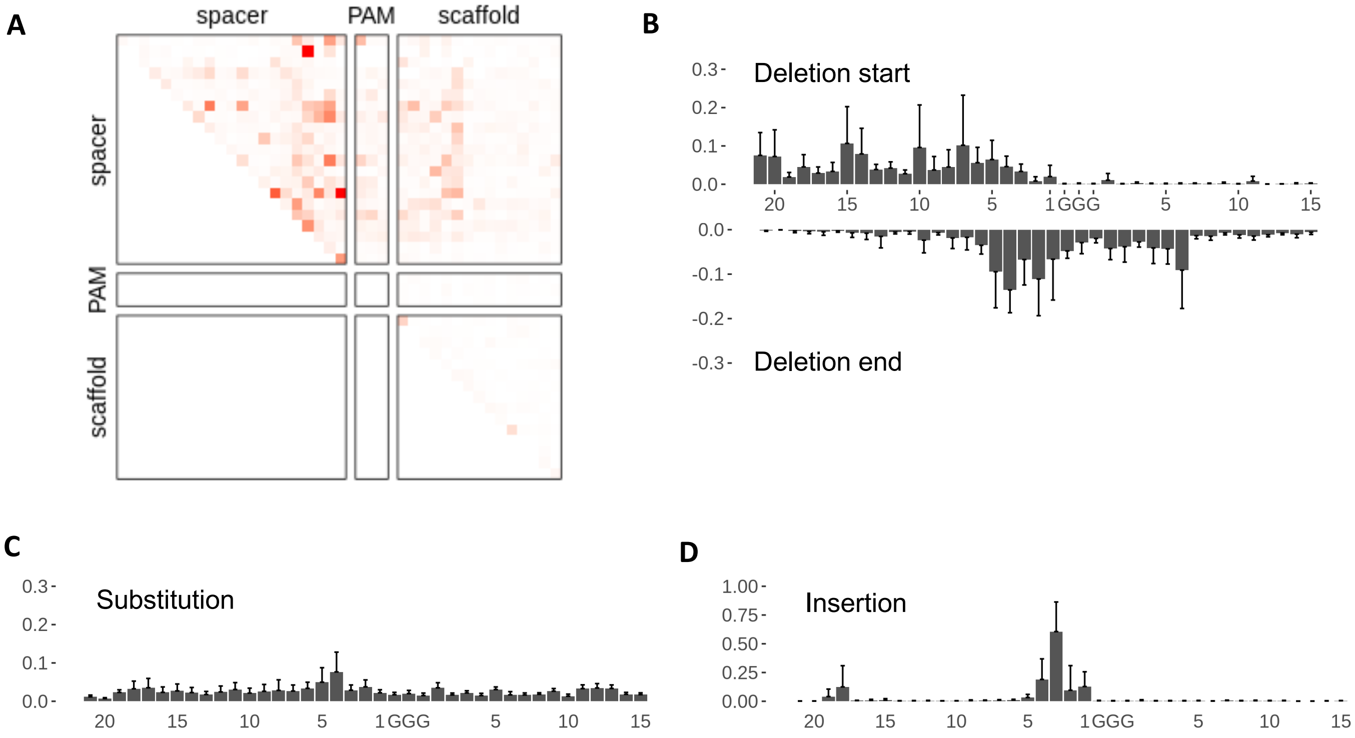


**Figure S9** The frequency of hgRNA-invitro A21 barcode's mutation position. **A** The heatmap showing the start and end of deletion. **B** The histogram of GESTALT's deletion position. **C** Substitution. **D** Insertion.

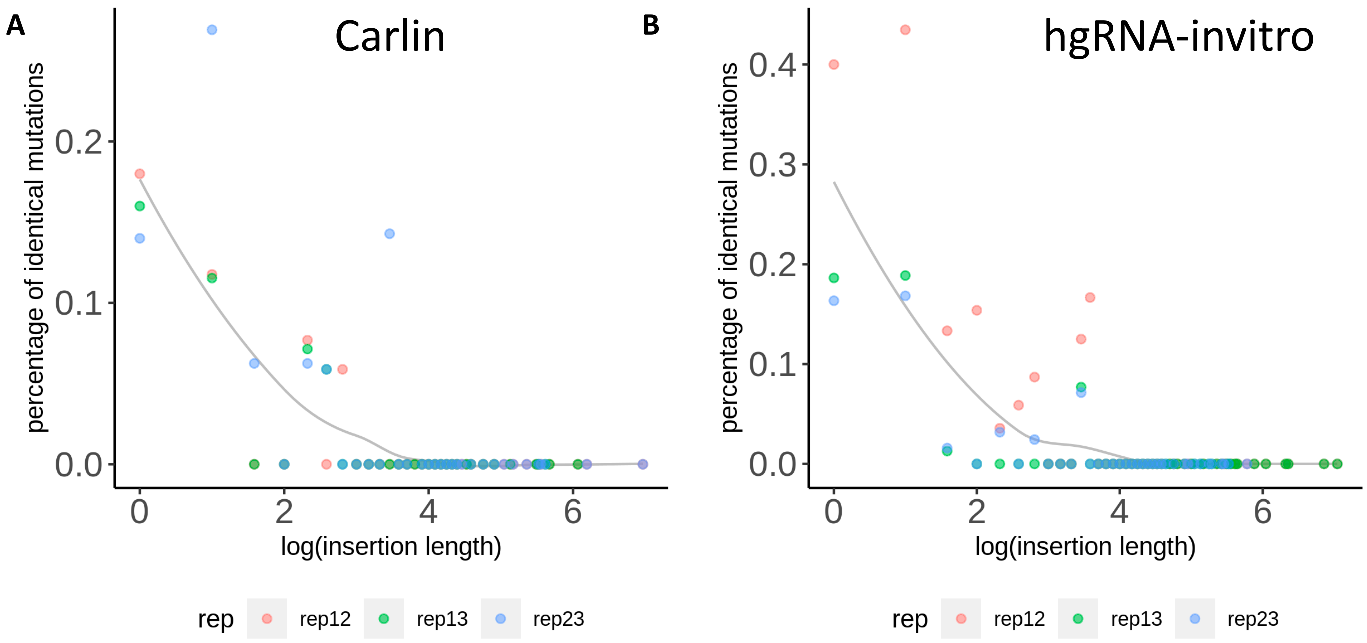


**Figure S10** The percentage of idenitical insertions between replicates. **A** The three replicates are low, medium, and high Dox at 96h of Carlin. **B** The three replicates are A'21-14d, A21-5d, and A21 pop6 of hgRNA-invitro.

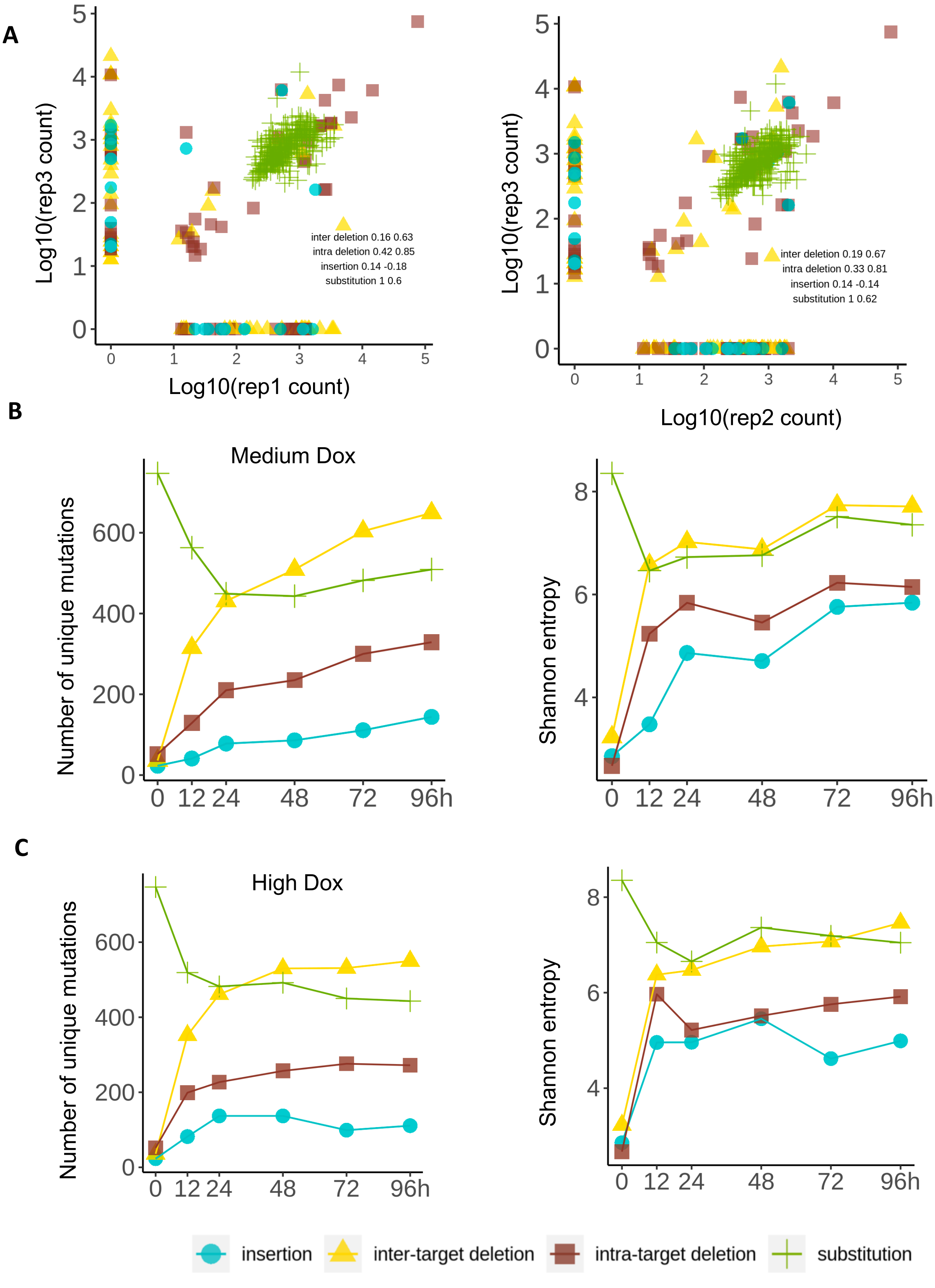


**Figure S11** The mutation dependency analysis of Carlin. **A** The percentage of overlapped mutations between replicates. The three replicates are negative Dox trial 1, negative Dox trial 2, and positive Dox at 0h. The text in the figure are overlapped percentage (how many points are in the diagonal) and the Pearson correlation of overlapped mutations (Pearson of diagonal points). **B-C** The Carlin's time-series data show changes of each mutation types through time.

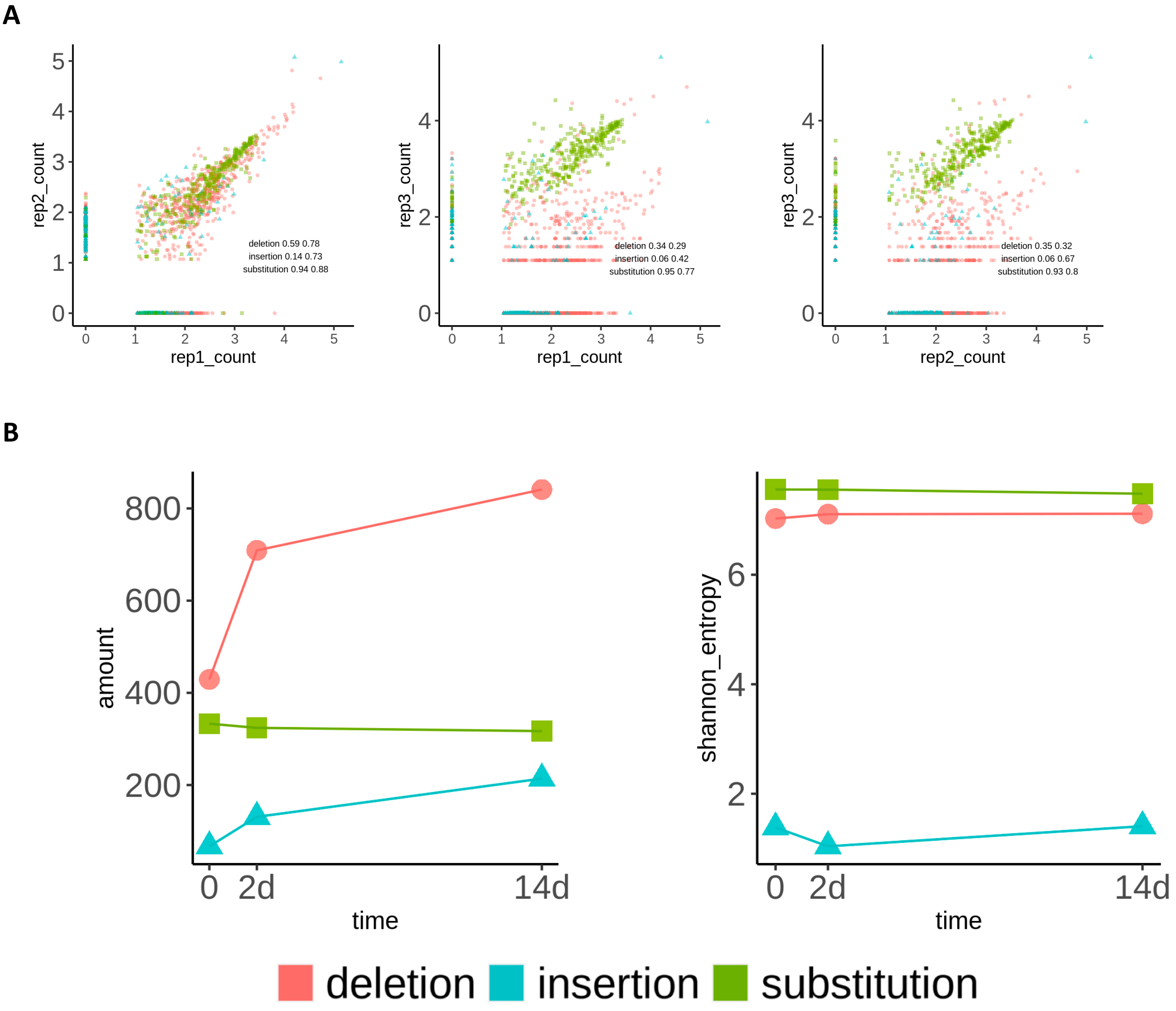


**Figure S12** The mutation dependency analysis of hgRNA-invitro. **A** The percentage of overlapped mutations between replicates. The three replicates are negative A'21-14d, negative A21-5d, and A21-pop6. The text in the figure are overlapped percentage (how many points are in the diagonal) and the Pearson correlation of overlapped mutations (Pearson of diagonal points). **B-C** The hgRNA-invitro's time-series data show changes of each mutation types through time.

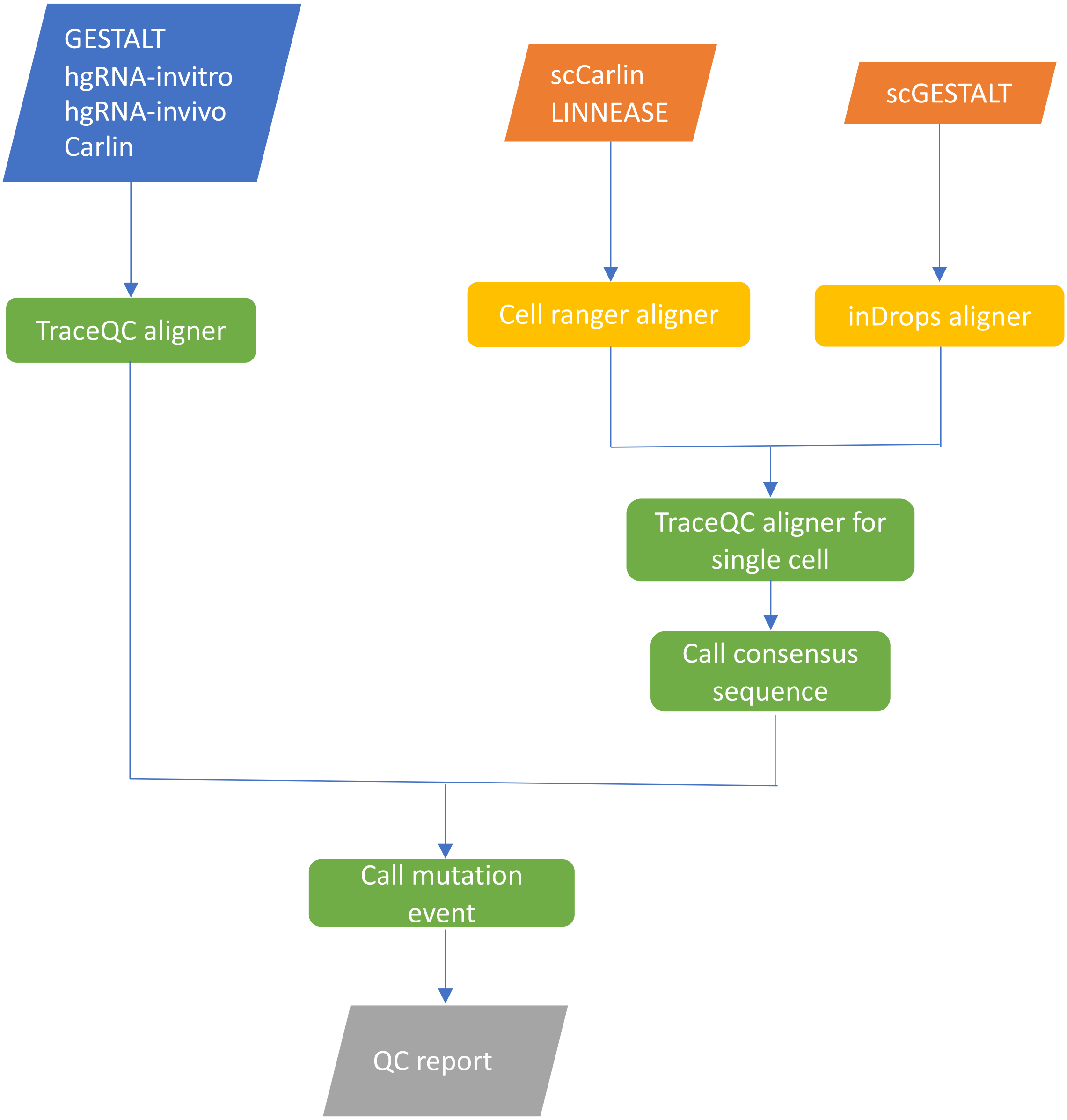


**Figure S13** The workflow of TraceQC processing pipeline in this study

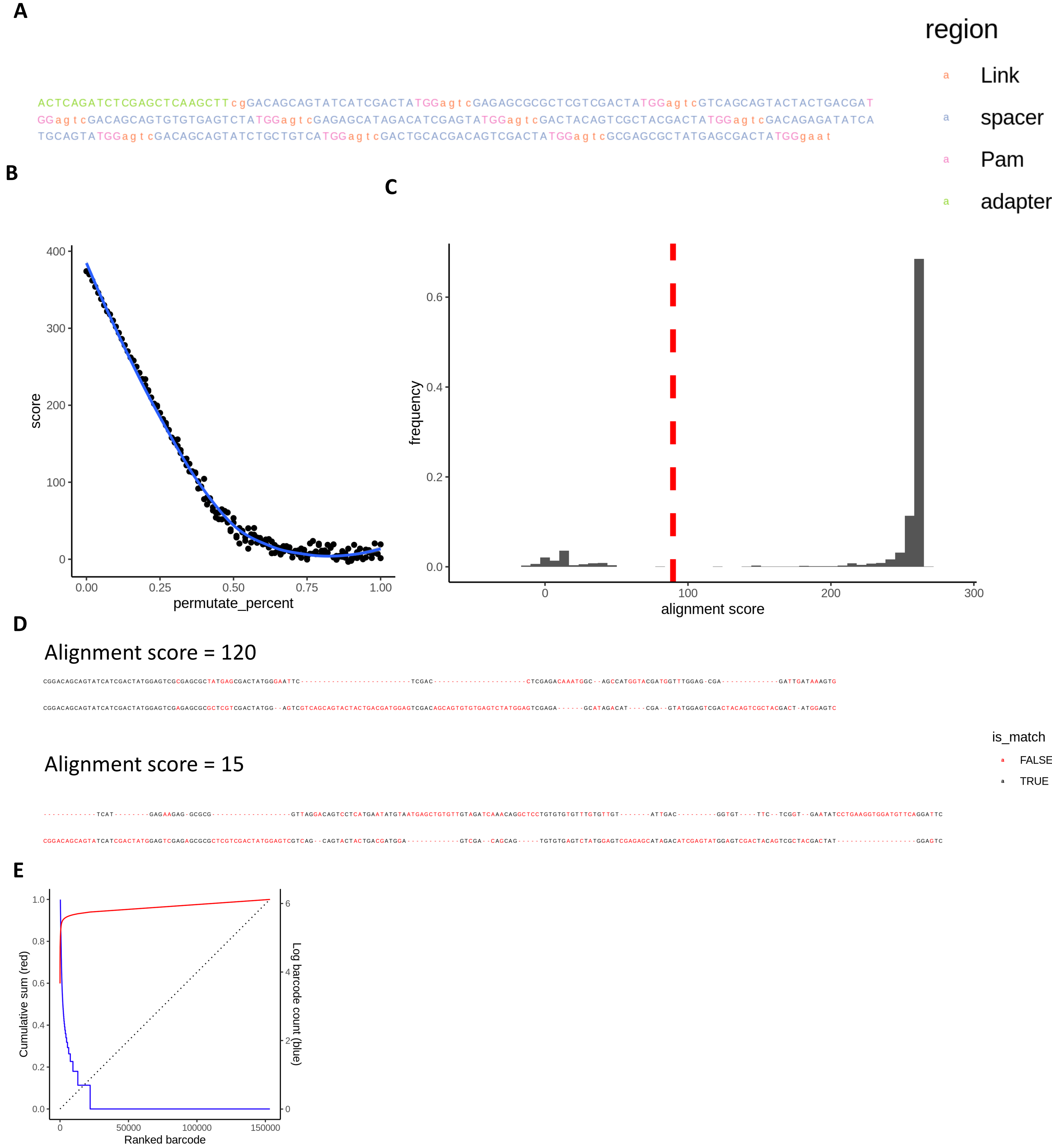


**Figure S14** Sequencing alignment result of GESTALT read 1. **A** Annotated GESTALT. v6 construct sequence. **B** Regression between alignment score and random permutated sequence percentage. **C** Histogram of alignment score of sample SRR3561150's R1 sequence. **D** Example alignment result of a good sequence and a bad sequence. The bad sequence are filtered out using selected permutate percentage as threshold. **E** Count distribution of sequences.

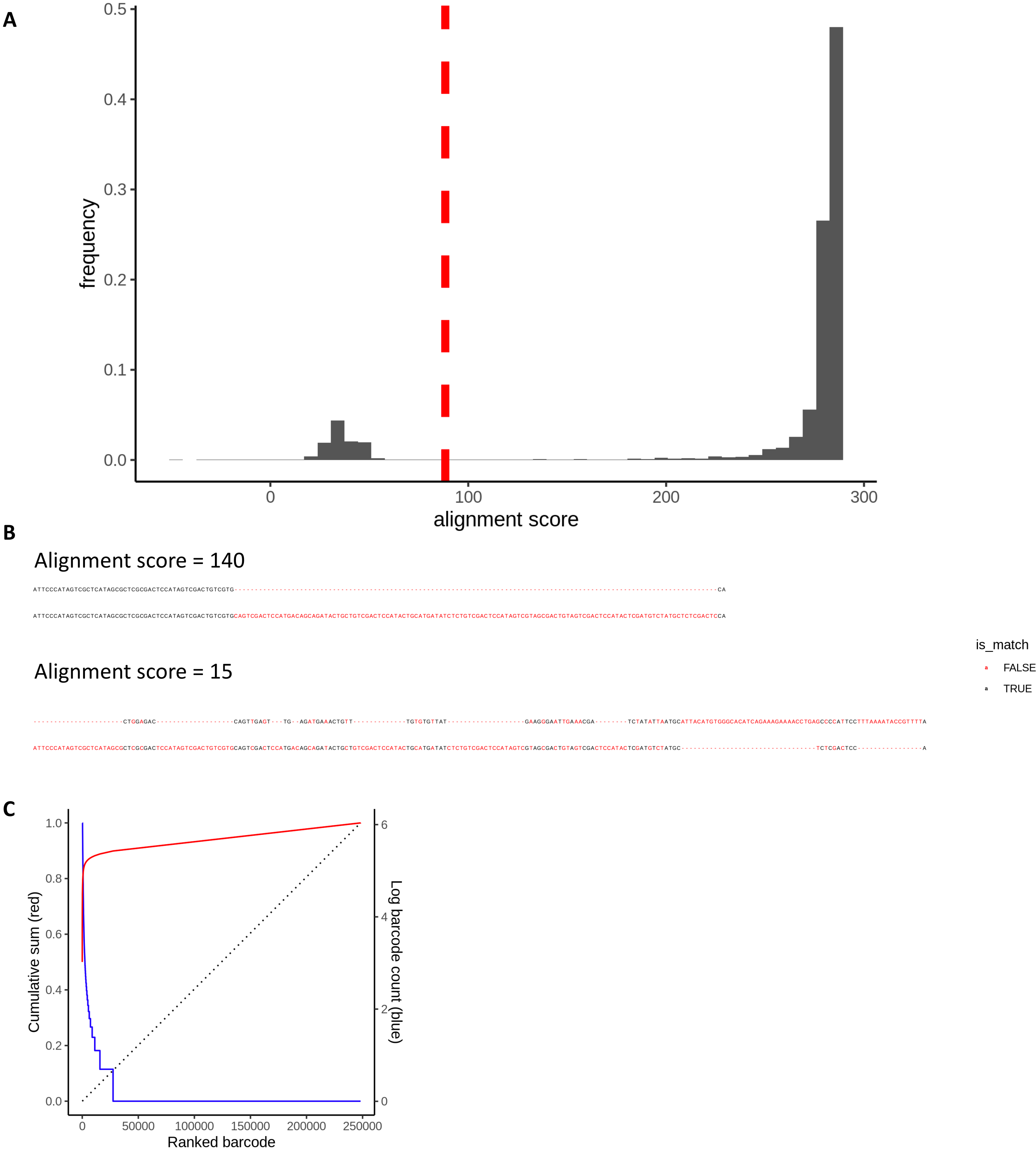


**Figure S15** Sequencing alignment result of GESTALT read 2. **A** Histogram of alignment score of sample SRR3561150's R1 sequence. **B** Example alignment result of a good sequence and a bad sequence. The bad sequence are filtered out using selected permutate percentage as threshold. **C** Count distribution of sequences.

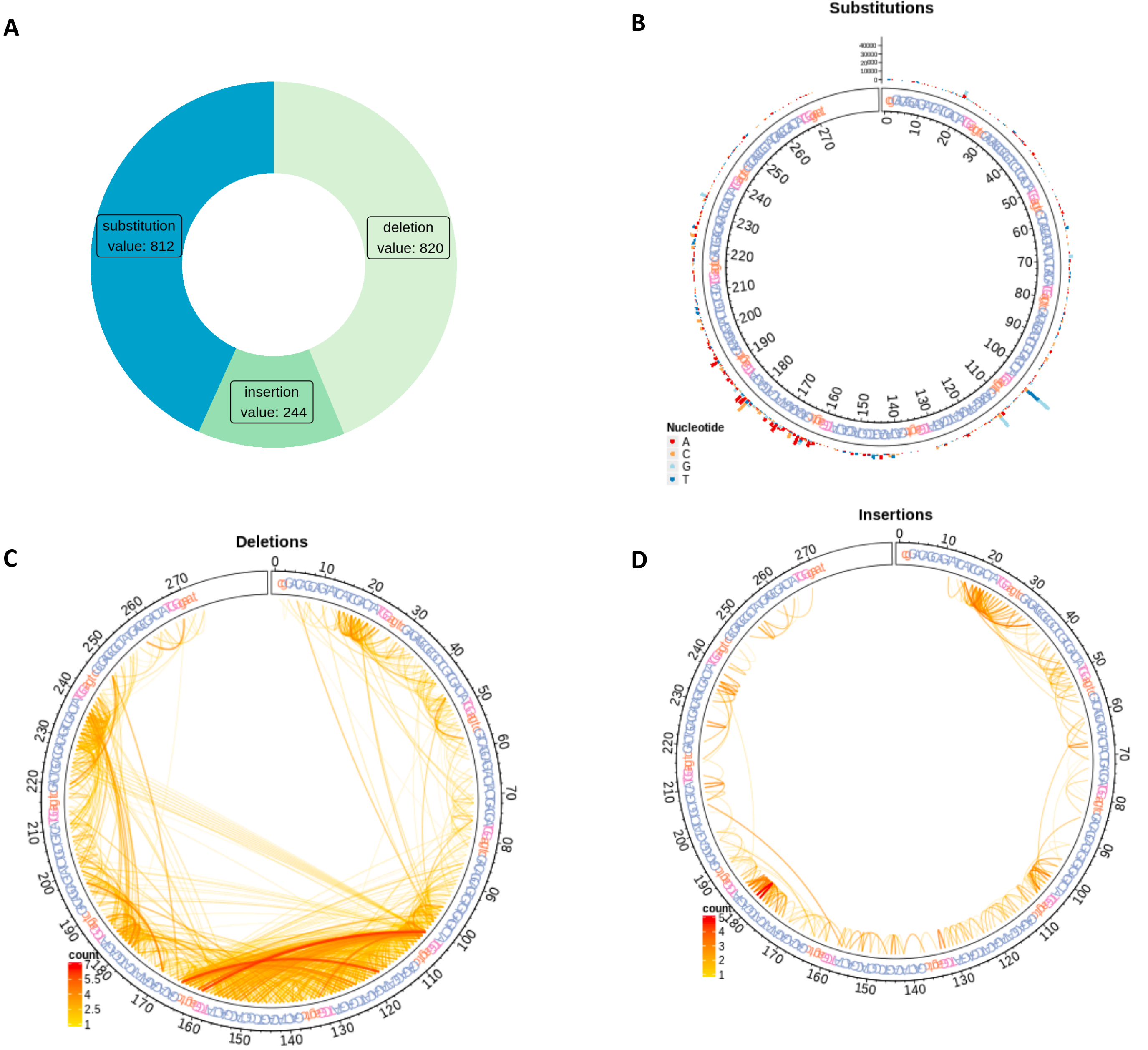


**Figure S16** Mutation characteristics of one sample from GESTALT: SRR3561150. **A** The percentage of three mutation types. **B** Substitution patterns of one barcode. **C** Deletion patterns. **D** Insertion patterns.

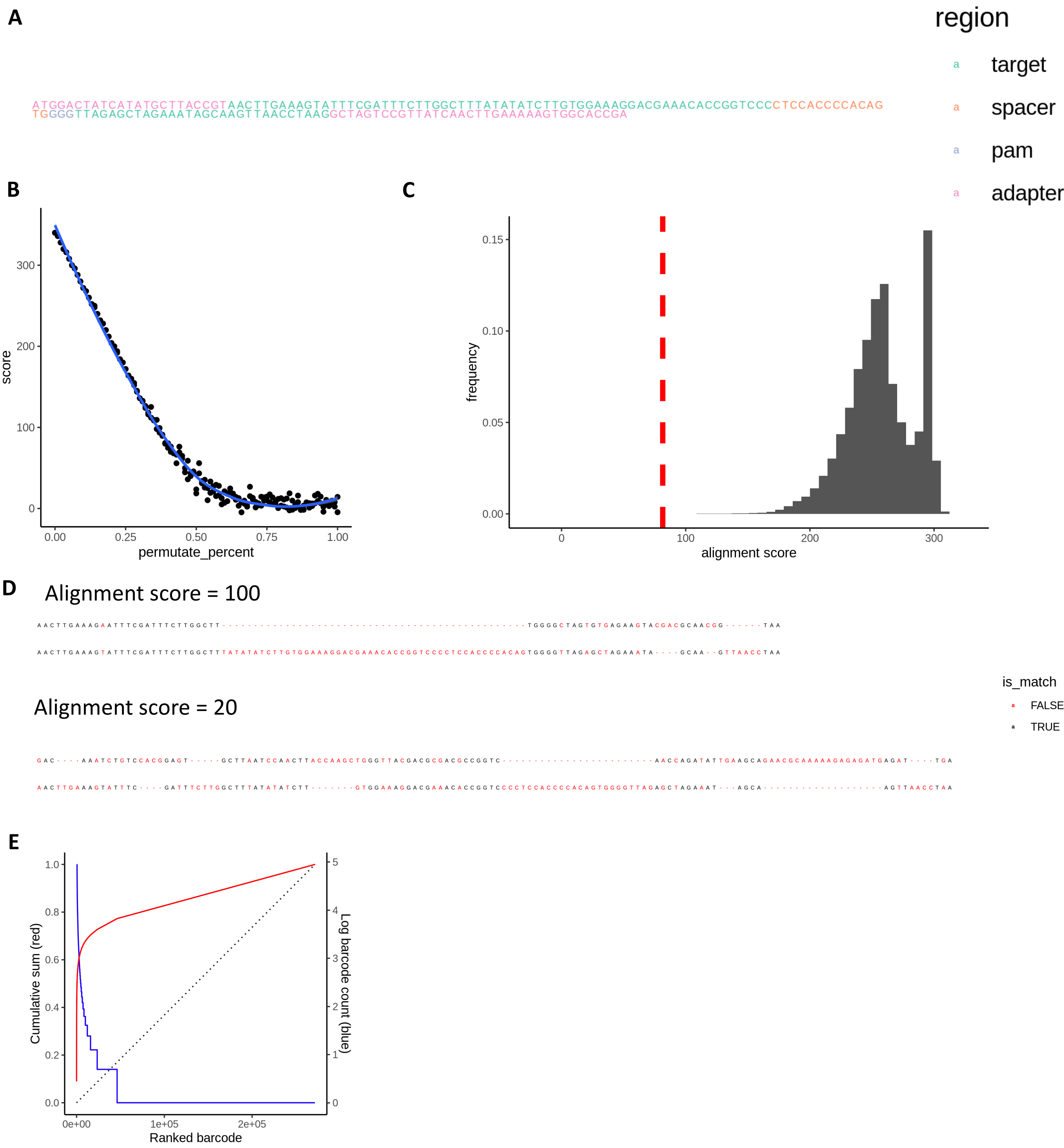


**Figure S17** Sequencing alignment result of hgRNA-invitro. **A** Annotated construct sequence. **B** Regression between alignment score and random permutated sequence percentage. **C** Histogram of alignment score of sample A21-14d. **D** Example alignment result of a good sequence and a bad sequence. The bad sequence are filtered out using selected permutate percentage as threshold. **E** Count distribution of sequences.

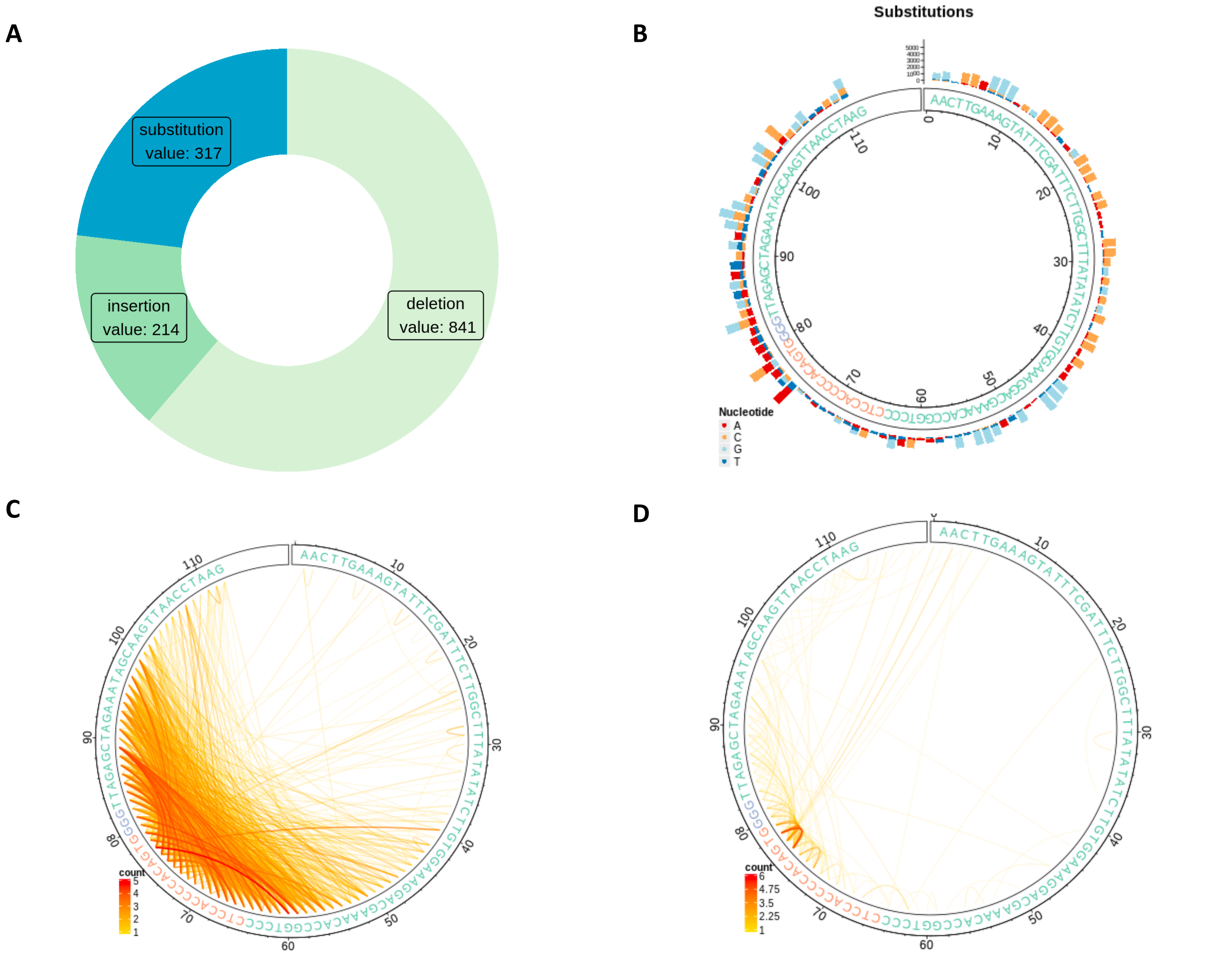


**Figure S18** Mutation characteristics of one sample from hgRNA-invitro: A21-14d. **A** The percentage of three mutation types. **B** Substitution patterns. **C** Deletion patterns. **D** Insertion patterns.

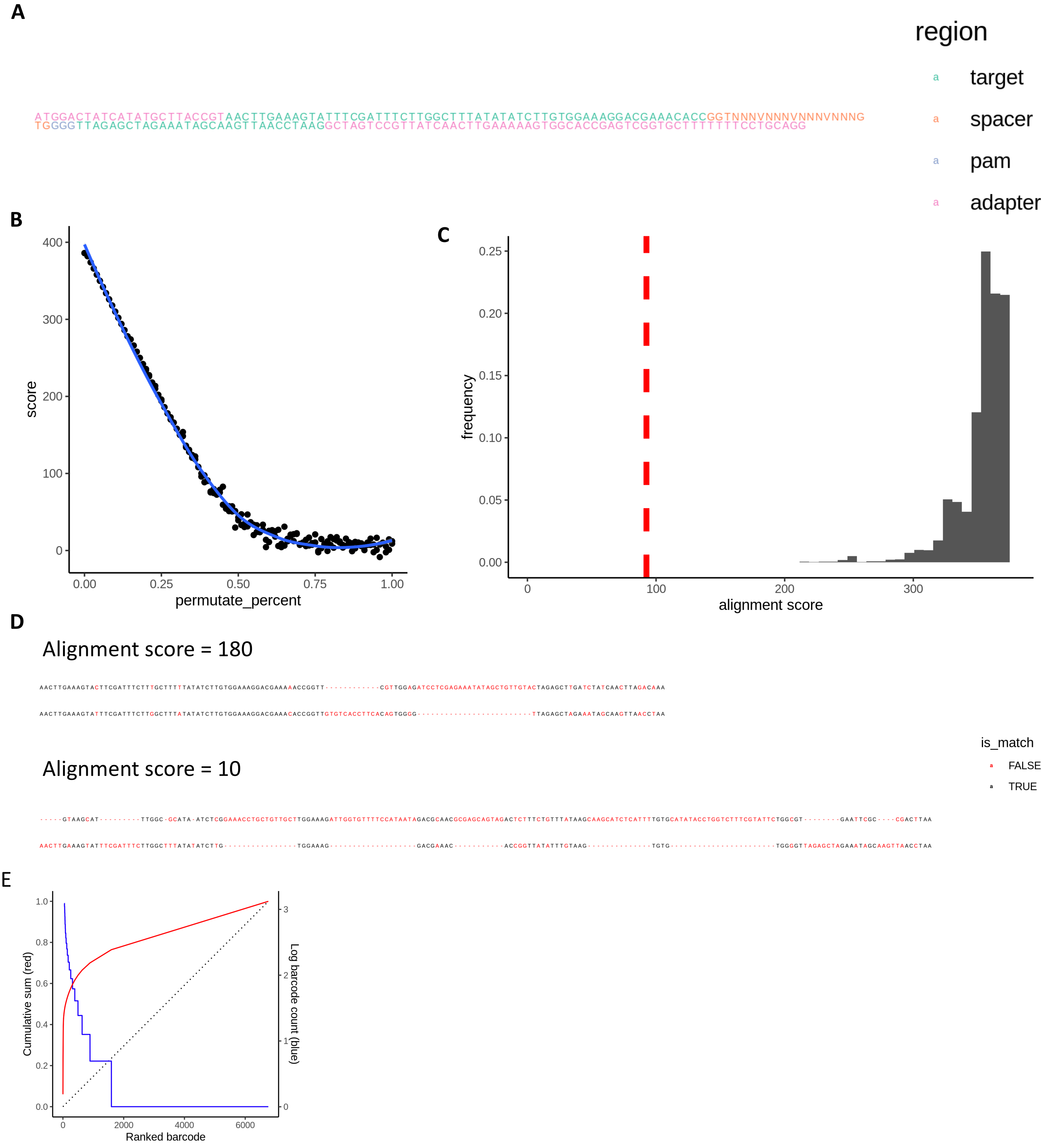


**Figure S19** Sequencing alignment result of hgRNA-invivo. **A** Annotated L21 construct sequence. **B** Regression between alignment score and random permutated sequence percentage. **C** Histogram of alignment score of sample A21-14d. **D** Example alignment result of a good sequence and a bad sequence. The bad sequence are filtered out using selected permutate percentage as threshold. **E** Count distribution of sequences.

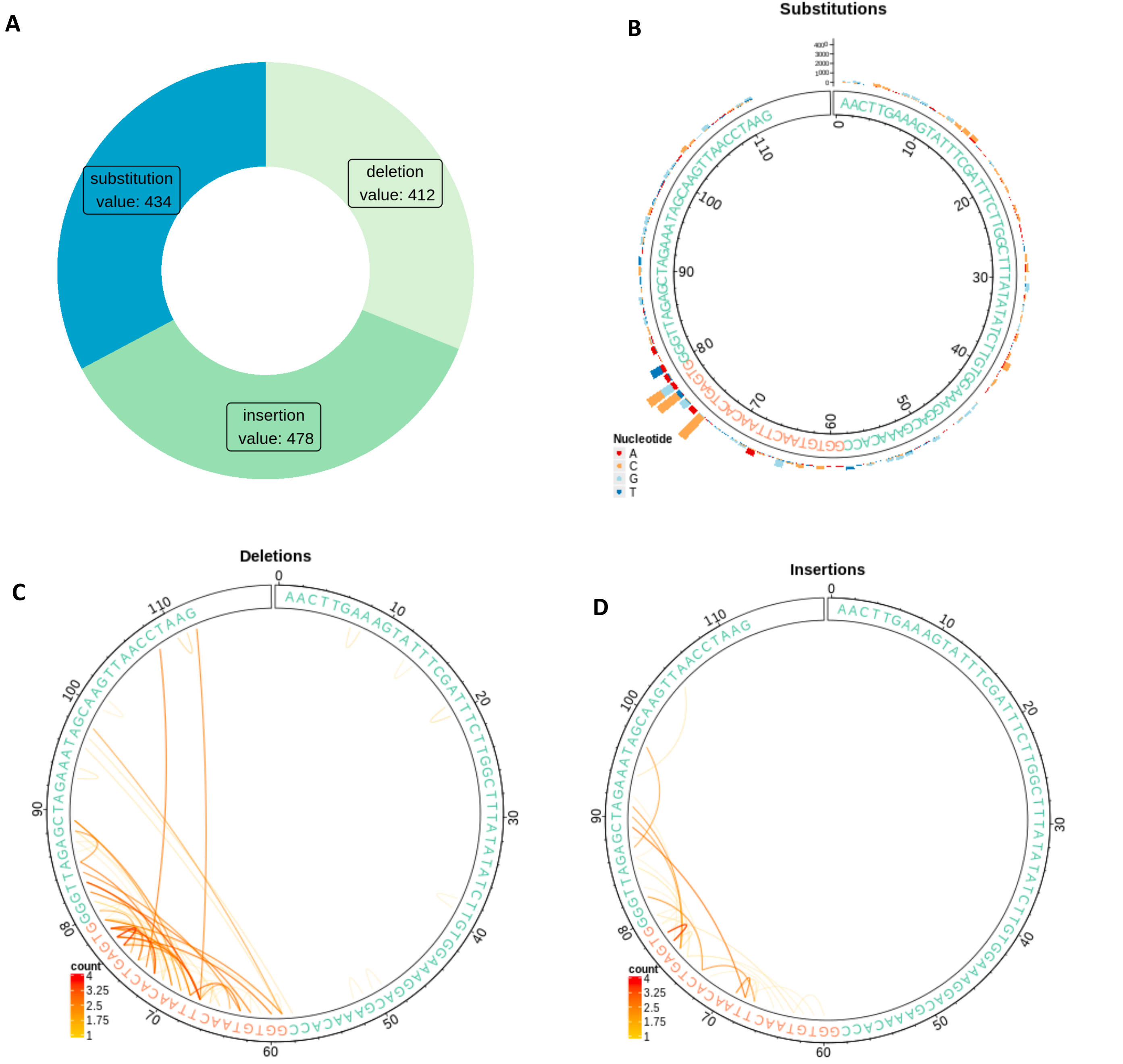


**Figure S20** Mutation characteristics of one selected sample of hgRNA-invivo: SRR7633623. **A** The percentage of three mutation types. **B** Substitution patterns of one barcode. **C** Deletion patterns. **D** Insertion patterns.

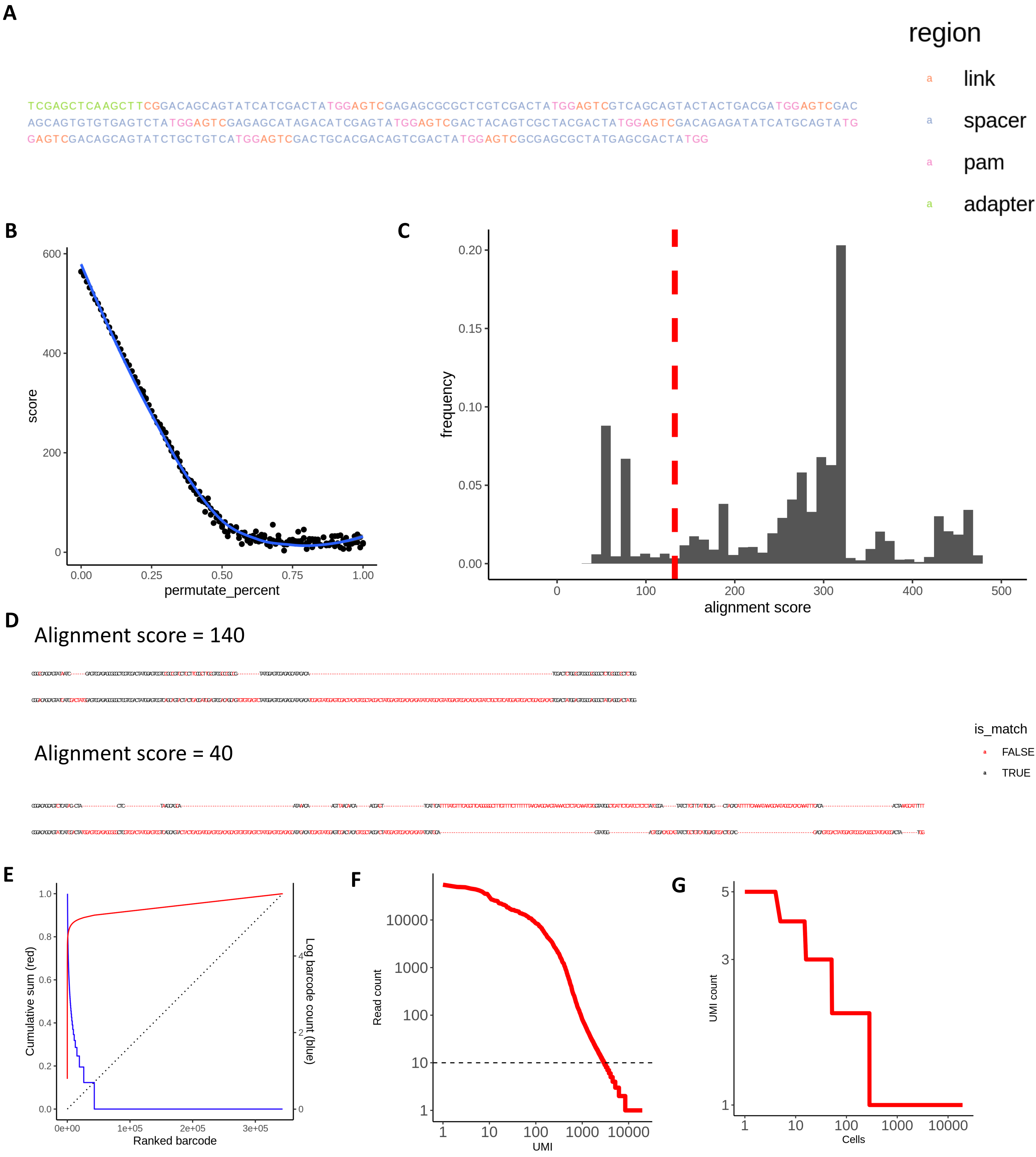


**Figure S21** Sequencing alignment result of scGESTALT. **A** Annotated construct sequence of scGESTALT. **B** Regression between alignment score and random permutated sequence percentage. **C** Histogram of alignment score of sample SRR6176748. **D** Example alignment result of a good sequence and a bad sequence. The bad sequence are filtered out using selected permutate percentage as threshold. **E** Count distribution of sequences.

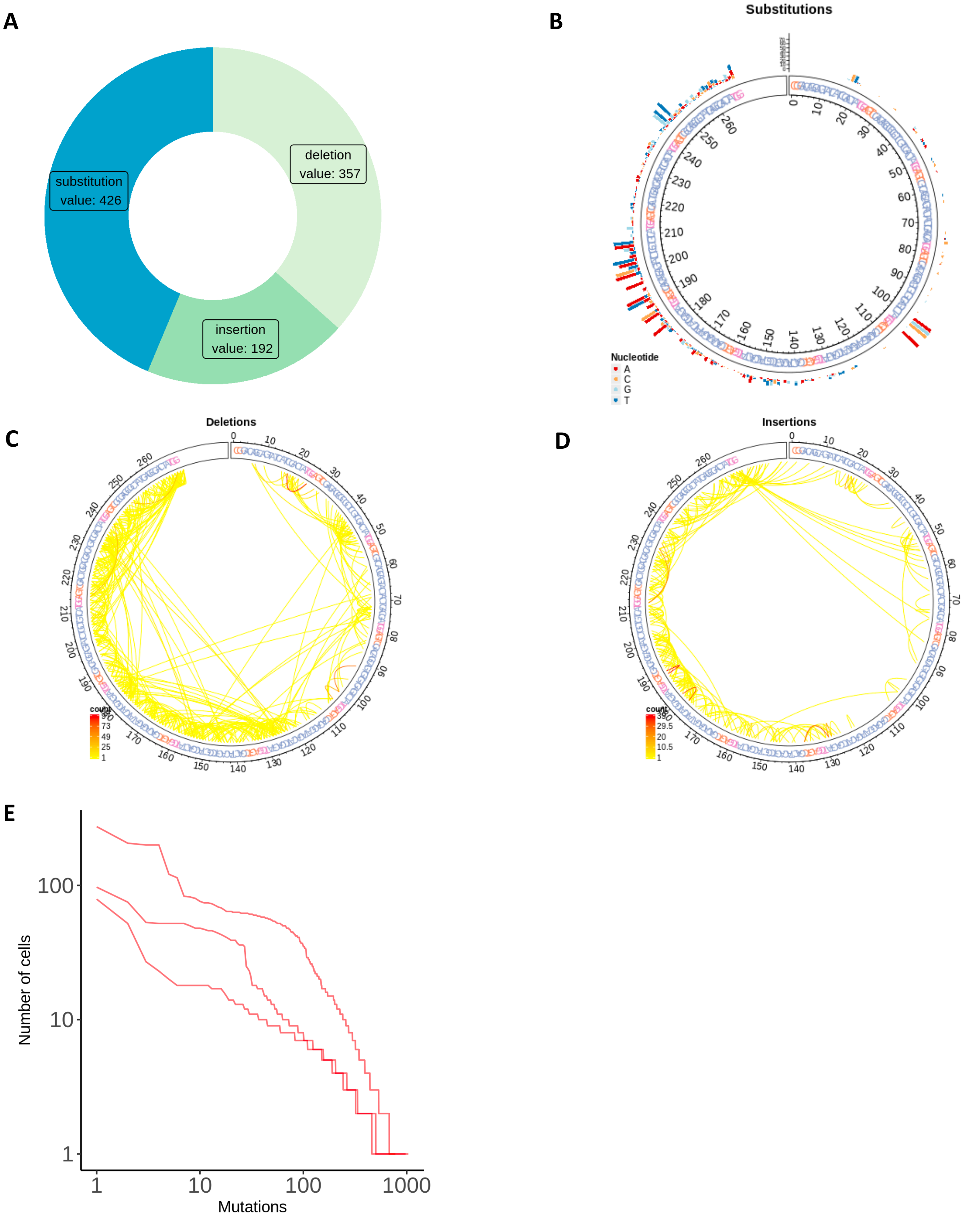


**Figure S22** Mutation characteristics of one sample of scGESTALT: SRR6176748. **A** The percentage of three mutation types. **B** Substitution patterns. **C** Deletion patterns. **D** Insertion patterns. **E** Mutation distribution of all single-cell samples.

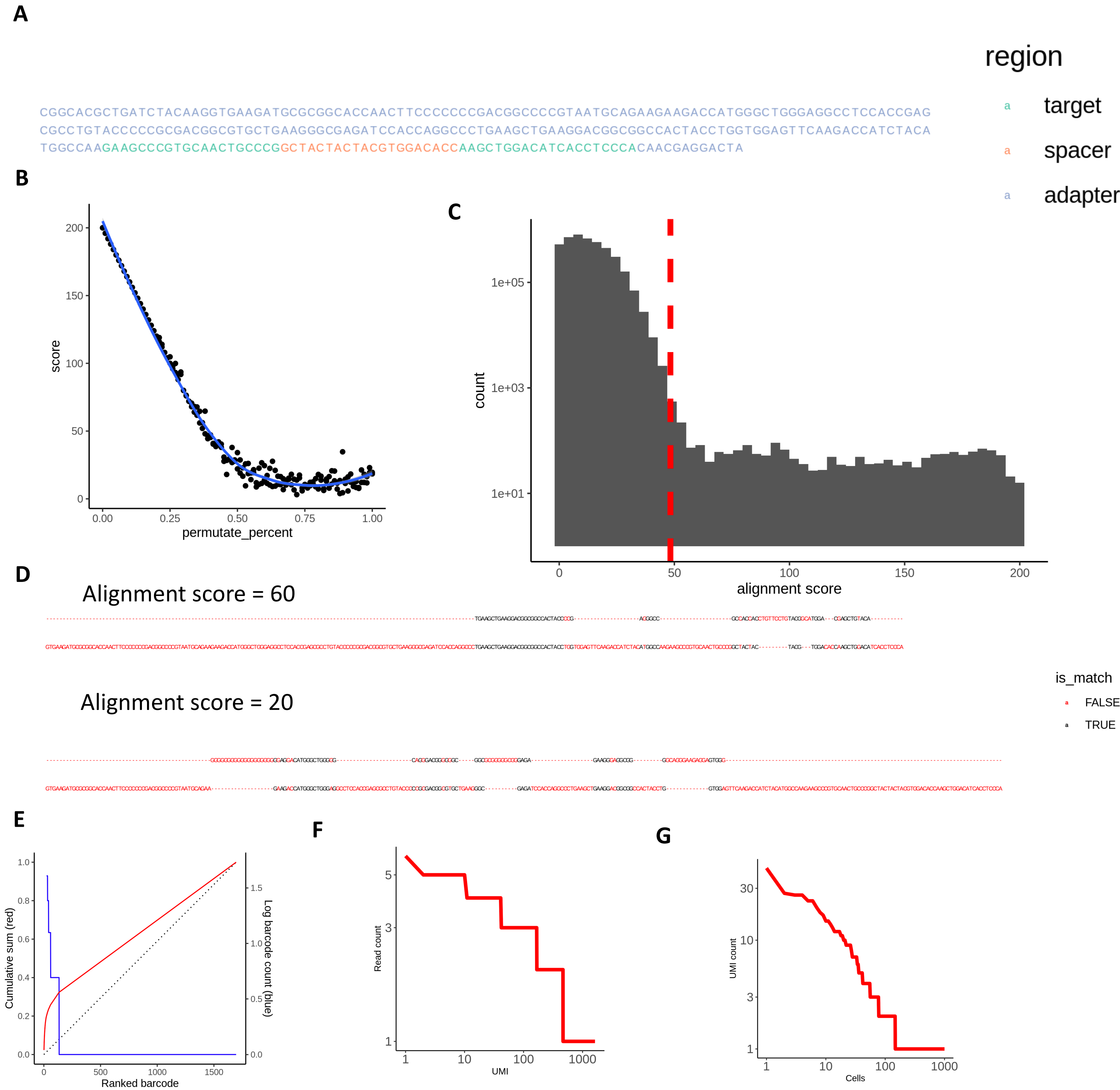


**Figure S23** Sequencing alignment result of LINNAESE. **A** Annotated construct sequence of LINNAESE. **B** Regression between alignment score and random permutated sequence percentage. **C** Histogram of alignment score of sample SRR6211489. **D** Example alignment result of a good sequence and a bad sequence. The bad sequence are filtered out using selected permutate percentage as threshold. **E** Count distribution of sequences.

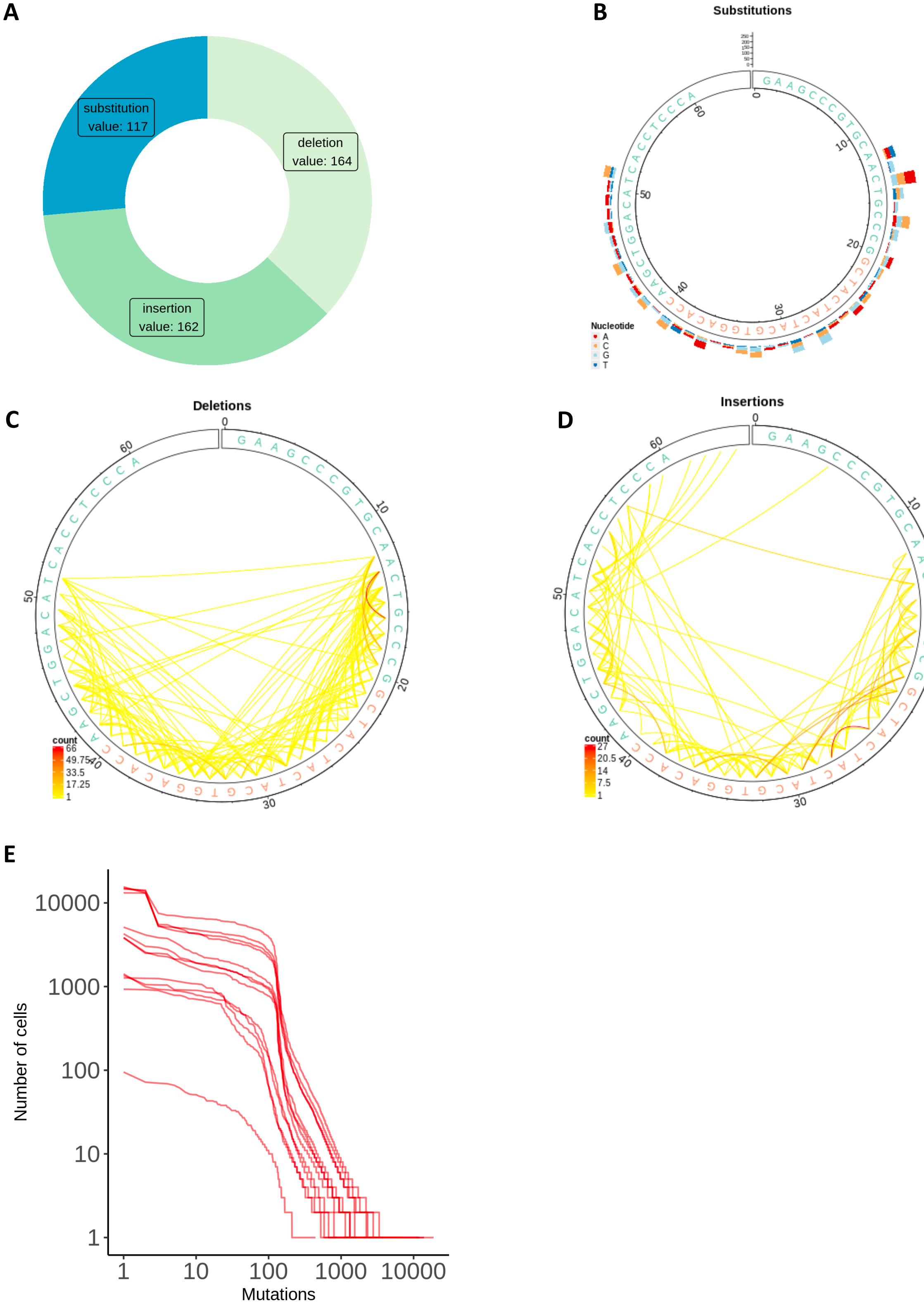


**Figure S24** Mutation characteristics of one sample of LINNAESE : SRR6211489. **A** The percentage of three mutation types. **B** Substitution patterns. **C** Deletion patterns. **D** Insertion patterns. **E** Mutation distribution of all single-cell samples.

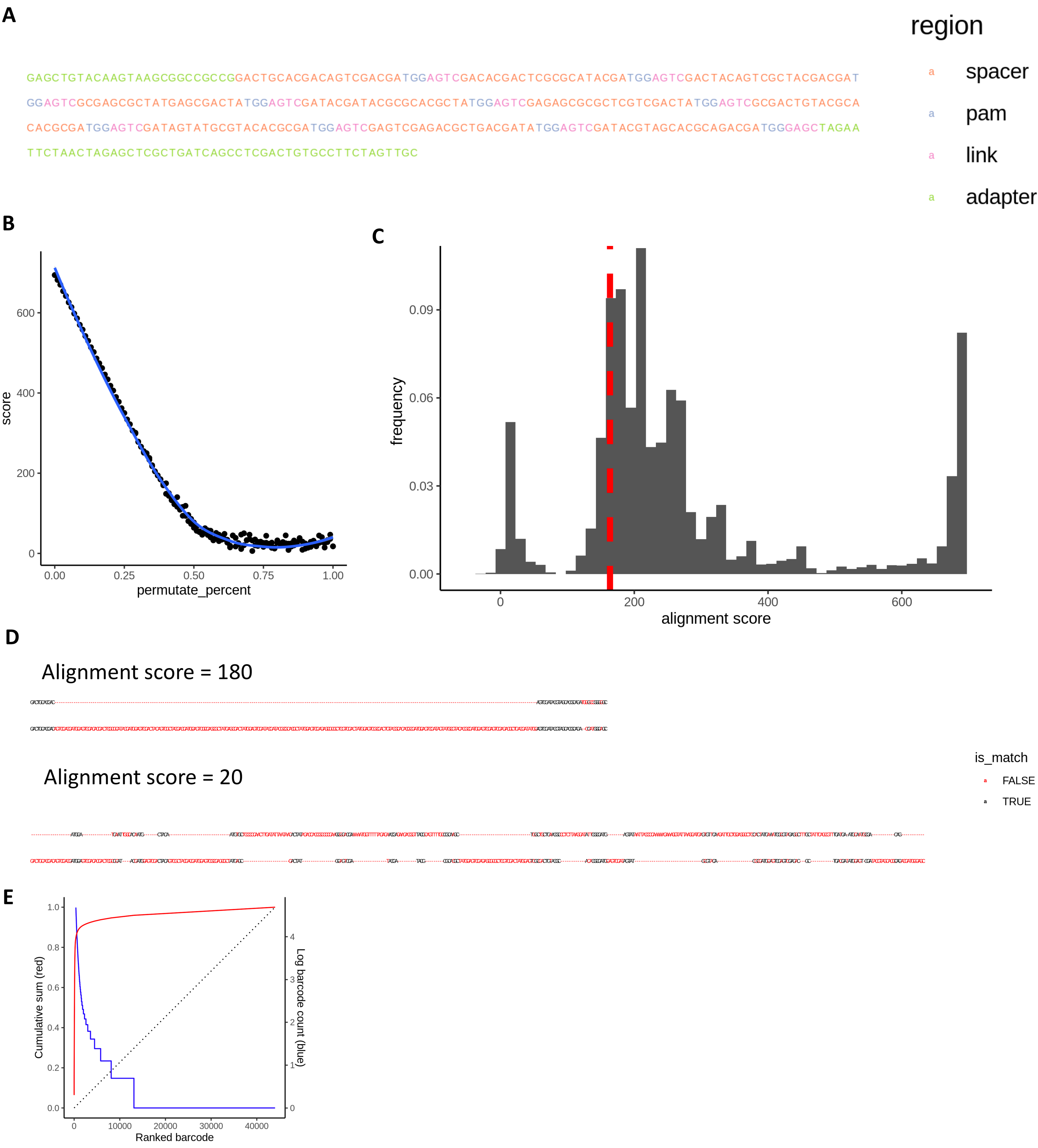


**Figure S25** Sequencing alignment result of Carlin. **A** Annotated construct sequence of Carlin. **B** Regression between alignment score and random permutated sequence percentage. **C** Histogram of alignment score of sample ChronicInduction_PosDox_Low_96h. **D** Example alignment result of a good sequence and a bad sequence. The bad sequence are filtered out using selected permutate percentage as threshold. **E** Count distribution of sequences.

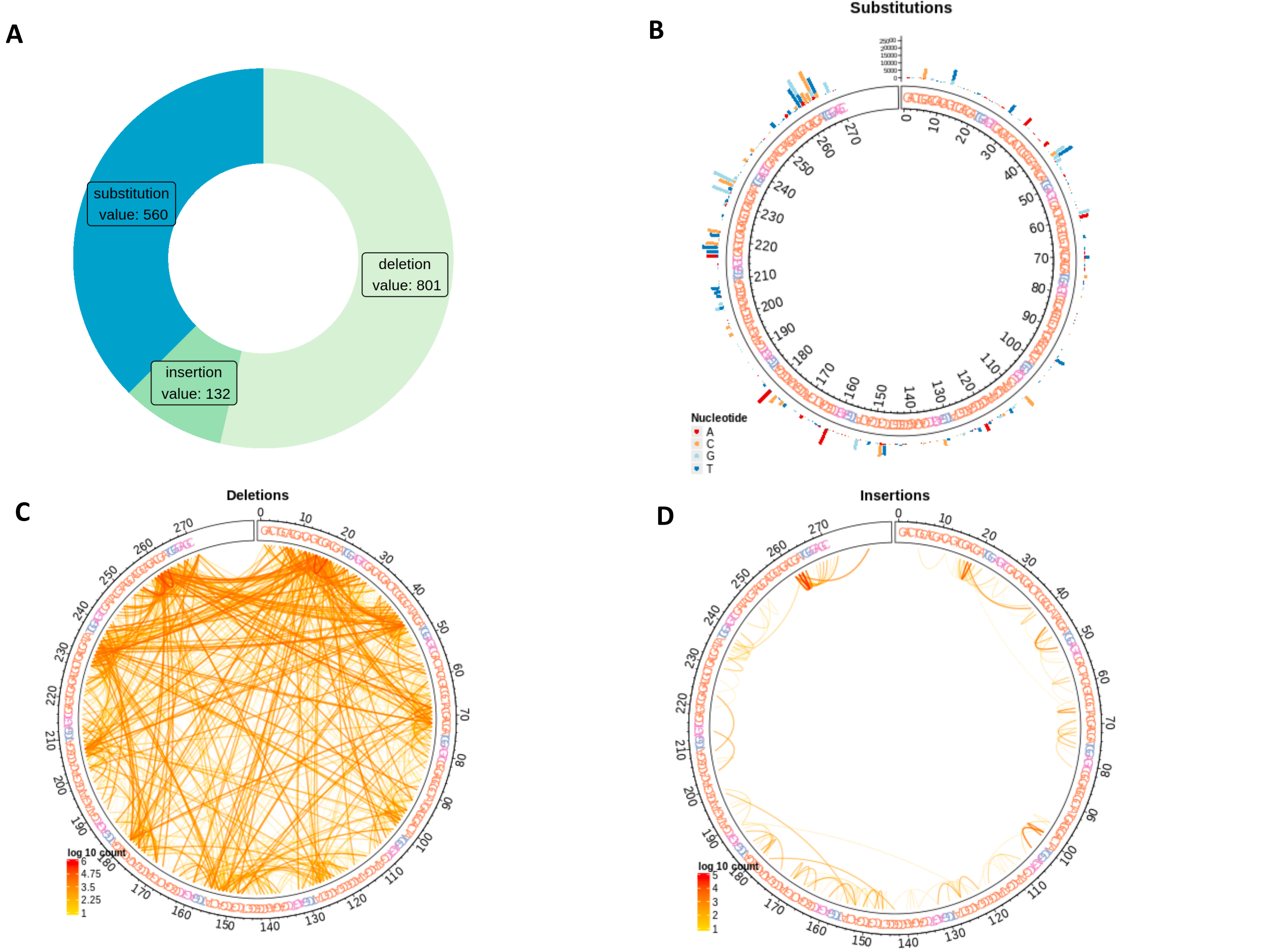


**Figure S26** Mutation characteristics of one Carlin sample: ChronicInduction_PosDox_Low_96h. **A** The percentage of three mutation types. **B** Substitution patterns. **C** Deletion patterns. **D** Insertion patterns. **E** The change of mutated percentage and Shannon entropy over time.

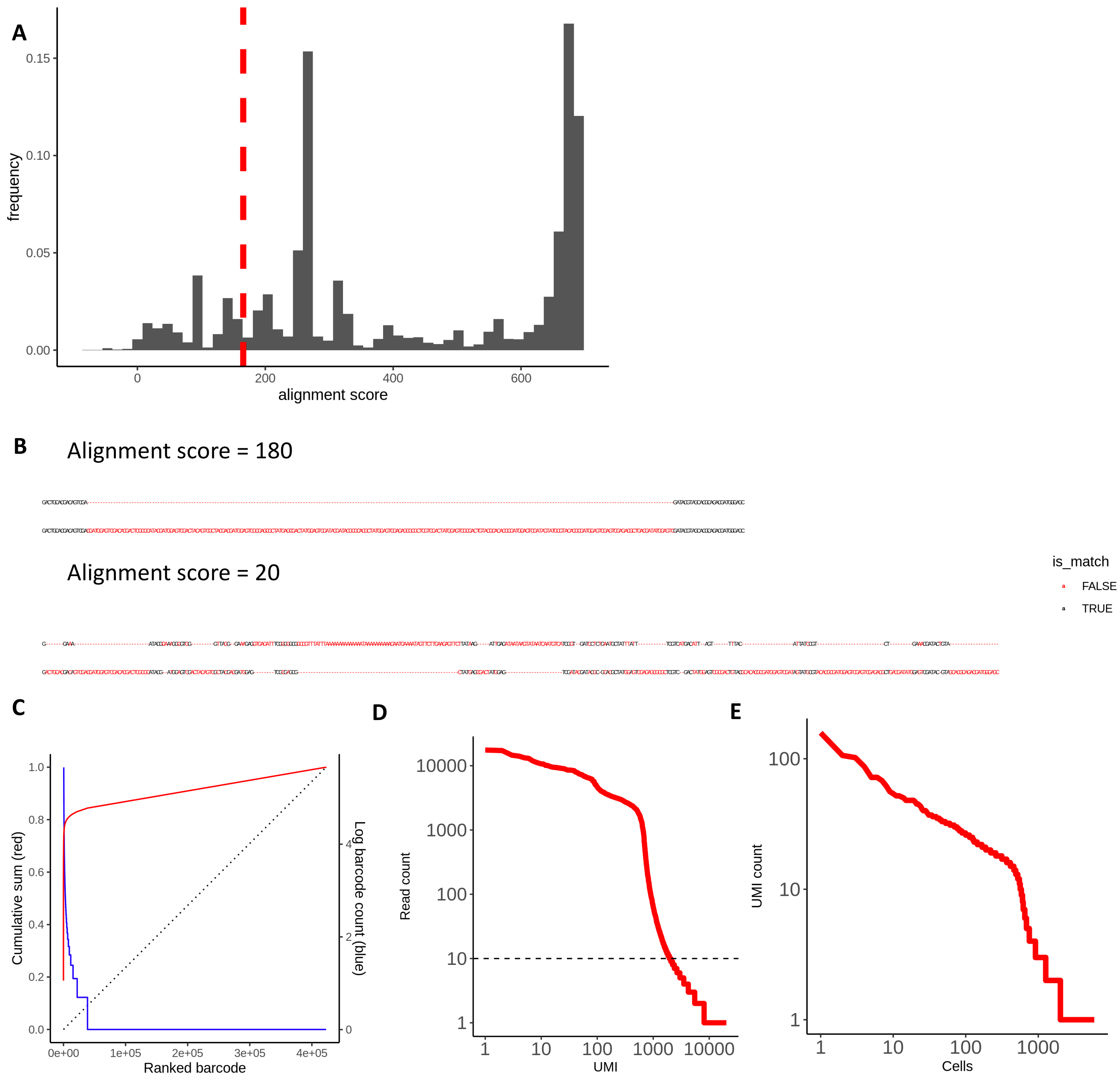


**Figure S27** Sequencing alignment result of scCarlin. **A** Histogram of alignment score of sample 5FU_FO817_SC. **B** Example alignment result of a good sequence and a bad sequence. The bad sequence are filtered out using selected permutate percentage as threshold. **C** Count distribution of sequences. **D** Read count per UMI. **E** UMI count per cell.

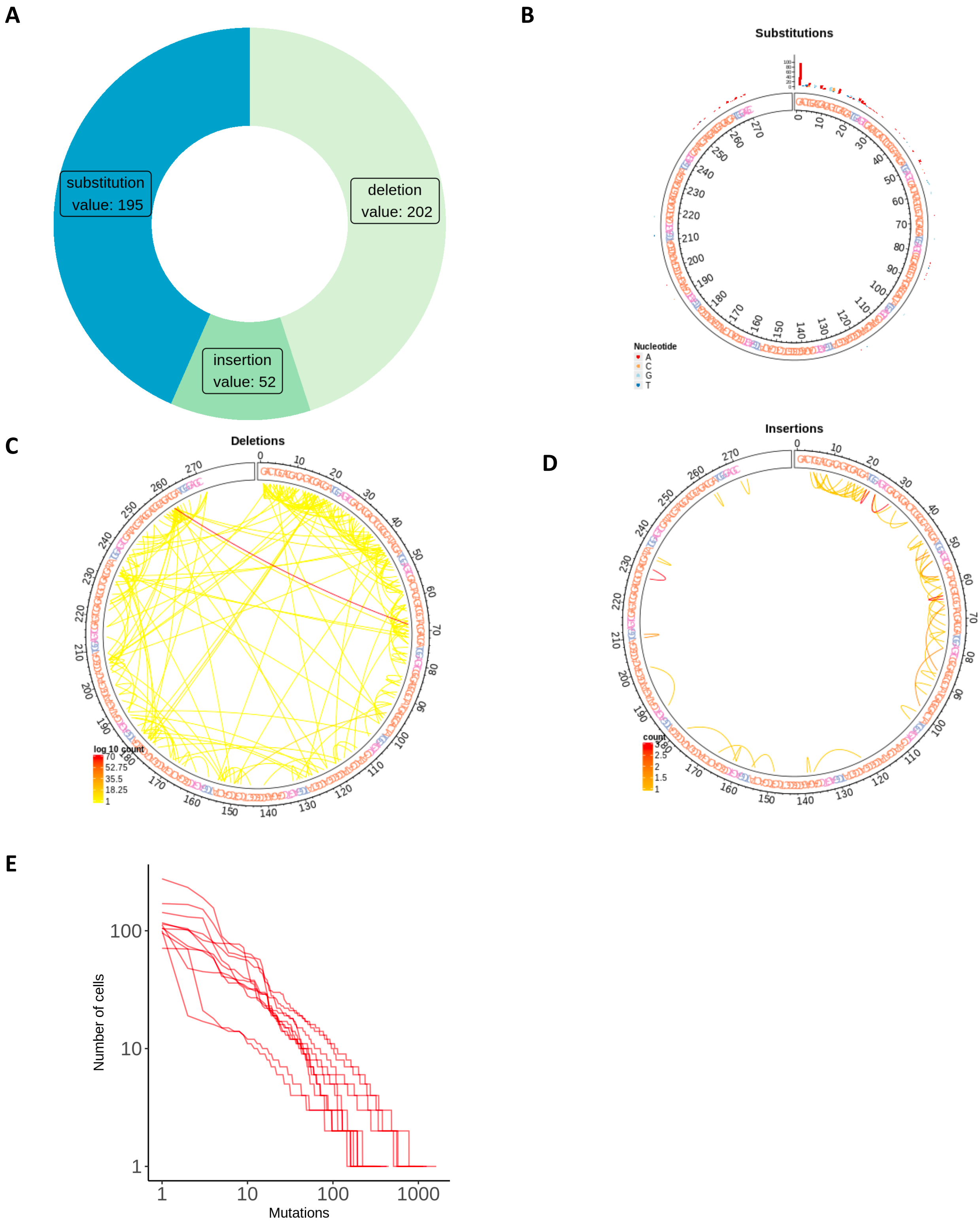


**Figure S28** Mutation characteristics of one sample of scCarlin: ChronicInduction_PosDox_Low_96h. **A** The percentage of three mutation types. **B** Substitution patterns. **C** Deletion patterns. **D** Insertion patterns. **E** Mutation distribution of all single-cell samples.
